# SALL1 and neural microenvironmental context specify identity of stem cell derived human microglia-like cells

**DOI:** 10.64898/2026.09.25.754357

**Authors:** Marina T. Nikolova, Kiara Freitag, Ryoko Okamoto, Andrea Catellani, Jennifer Hinke, Nicolas Luginbühl, Hsiu-Chuan Lin, Johanna Kaindl, Zhisong He, Philipp Wahle, Juliane Rohland, Laufey Geirsdottir, J. Gray Camp, Beate Winner, Ido Amit, Barbara Treutlein

## Abstract

Microglia, the immune cells of the brain parenchyma, play a pivotal role in neurodevelopment and neuroprotection. Human stem cell-derived microglia-like cells (MGL) offer a valuable, human-relevant system to study microglia in vitro. However, their cell states and intercellular interactions within complex 3-dimensional (3D) tissue contexts in vitro remain underexplored. Here, we analyze the gene expression profiles of MGL generated by two differentiation protocols in conventional 2D cultures. We show that co-culture with neural organoids promotes a more mature, in vivo–like MGL phenotype but this signature is lost during long-term co-culture. By varying media compositions, we modulate 2D MGL states toward more active or more homeostatic identities, which may enhance the similarity to primary microglia identities upon subsequent co-culture. Moreover, forced expression of the microglial lineage regulator SALL1 shifts MGL away from a general myeloid state toward a more defined pre-microglial fate. SALL1 overexpression within a 3D neural organoid environment further amplifies the shift toward a homeostatic developmental microglia population. Collectively, our transcriptomic analysis delineates human MGL states in 2D and 3D contexts, refines existing differentiation protocols toward more authentic MGL, and establishes a framework to investigate the intricate microglia–neural tissue interplay.

## Introduction

Microglia are the resident immune cells of the brain parenchyma and act as gatekeepers of the central nervous system (CNS) (see Table S1 for all abbreviations) ^1,2^. They originate from erythro-myeloid progenitors in the yolk sac early in development ^3^ and appear in the CNS as early as the 5th gestational week in humans ^4^ or embryonic day 9 in mice ^3^. Once having populated the CNS, microglia self-renew locally and maintain a constant pool of cells ^5^, constituting 5-15% of all cells in the CNS, depending on brain region and health status ^6^. They play a key role in the proper development and in maintaining the homeostasis of the CNS via synaptic pruning, pathogen recognition, cytokine release, secretion of neurotrophic factors, and phagocytizing apoptotic cells and cellular debris, among others ^7^. Impairment in any of these key functions and in microglia interaction with other cells in the CNS has been linked to the occurrence of neurodegenerative and neurodevelopmental disorders, such as Alzheimer’s disease ^8^ and Autism spectrum disorder ^9^, respectively.

During their development, microglia in the white matter undergo a transient phase, known as proliferative-region-associated microglia (PAM) 10. PAM in mice express a core signature including Spp1, Gpnmb, Clec7a and Itgax, shared with disease-associated microglia (DAM) ^10^, which are enriched in aging and neurodegenerative conditions ^11^. In contrast, border-associated macrophages (BAMs) populate CNS interfaces such as the meninges, perivascular spaces, and choroid plexus, forming a long-lived immune barrier distinct from parenchymal microglia ^12^. While PAMs represent a developmentally programmed, transient microglial phenotype, BAMs maintain specialized roles in immune surveillance and barrier regulation throughout life. Microglia research has traditionally relied on primary cells and animal models, but these are not without drawbacks. Access to human microglia from primary tissues is limited, with cell phenotypes often being obscured by artifacts resulting from ante-mortem conditions, post-mortem delay or tissue processing. Furthermore, primary microglia have been shown to undergo drastic transcriptional and functional changes, such as downregulation of the microglial core homeostatic markers TMEM119, P2RY12 and SALL1, after being removed from their native in vivo environment and cultured in a dish for as little as 6 hours ^13-15^. While animal models have been instrumental in uncovering the ontology and function of microglia, recent studies highlight essential differences between human and non-human animal microglia, including the expression of genes implicated in brain development and neurodegenerative diseases, as well as age-related changes, which could hamper translation of studies from different species into the clinic ^14,16^. Hence, there is a need for the establishment of robust and scalable protocols for microglia derivation in vitro, which recapitulate microglia states in vivo.

Recent breakthroughs in stem cell technologies have enabled the expandable generation of human induced pluripotent stem cells (hiPSCs) from various cell types and their differentiation into cell and tissue avatars of their in vivo counterparts ^17,17^. Multiple protocols for the generation of stem cell-derived microglia (microglia-like cells, MGL) have been published ^19^ and used to study progranulin-deficient frontotemporal dementia ^20^, Alzheimer’s disease ^21^, Huntington’s disease ^22^ and others. However, MGLs do not fully recapitulate all aspects of microglial characteristics such as comparable expression of TMEM119, P2RY12 or SALL1 ^23^, or microglia ontogeny ^24^. The transcription factor (TF) SALL1, in particular, has been identified as a key transcriptional regulator of microglia identity and phenotype, differentiating microglia from other macrophages ^14,25-27^. Studies suggest that Sall1 expression is essential for suppression of the BAM marker CD206 (encoded by Mrc1) ^28^. To address these issues, several studies have successfully transplanted human MGLs into mouse brains and shown increased similarity to microglia in vivo ^29-34^. However, in order to epitomize human CNS characteristics in a scalable manner, the coculture of human-only MGL cells with brain organoids has emerged as a promising approach ^35-38^. These co-cultures have so far been unable to reach the similarity to in vivo microglia achieved upon xenotransplantation, unless being transplanted into an animal host ^39^. These MGL typically lack mature microglia-specific genes and exhibit a phenotypic profile more similar to immature macrophages ^40,41^. Thus the challenge to generate reliable and specialized human microglia in vitro remains.

Here we present an in-depth single-cell transcriptomics characterization and optimization of MGL in 2D monoculture and in co-culture within human neural organoids. Our study highlights the high heterogeneity within MGL in vitro, as well as their increasing similarity to human primary developing microglia upon neural organoid co-culture. We uncover remaining differences between the primary and stem cell-derived microglia and further identify culture conditions for the generation of in vivo-like MGL in vitro. We overexpress the key microglia TF SALL1 in differentiating MGL and detect a decrease of yolk sac myeloid cell signature and increase of markers of early microglia cells. SALL1 over-expression in MGL co-culture with neural organoids also led to enhanced microglia signatures related to nervous system development. Thus, we provide a resource of various MGL states in 2D culture and upon co-culture with neural organoids and lay the groundwork for an optimized MGL protocol that generates microglia with an in vivo-like transcriptome and that can be successfully utilized for long-term co-culture with brain organoids.

## Results

### Human MGL exhibit diverse cell states which partially resemble human primary microglia

To generate MGLs for co-culture with neural organoids, we first explored two established protocols for iPSC to MGL differentiation for scalability, marker expression and resemblance to primary microglia (Protocol I see Haenseler et al. 19 and Brownjohn et al. ^42^, Protocol II see Lanfer et al. ^43^, (Fig. S1A, B). Both Protocols yielded MGL with comparable morphology and IBA1 expression and allowed weekly collection of MGL precursors for up to one month.

We performed single-cell RNA-sequencing (scRNA-seq, 10X Genomics) of MGL generated from both protocols using the same human embryonic stem cell line (day 9 of final differentiation, day 28 of culture; 6,814 and 5,577 cells, respectively) and identified largely overlapping transcriptomic profiles with similar proliferation signatures (Fig. S1C-E). Nearly all cells expressed core myeloid markers (AIF1/IBA1, TREM2, PTPRC, CSF1R), but clustering revealed notable heterogeneity (Fig. S1F-H): We observed a majority ‘homeostatic MGL’ population with sparse expression of homeostatic microglia markers (GPR183, P2RY12, P2RY13, CX3CR1, TMEM119) ^44^ and a CNS-associated macrophage signature (F13A1, STAB1, RNASE1), but no detectable SALL1 expression ^45^, a ‘remodeling MGL’ population enriched in extracellular matrix remodeling genes (MMP7, TIMP3), a minor ‘activated MGL’ population expressing HLA-DR genes, and several off-target clusters resembling neutrophils (RETN, CEACAM8), mast cells (CLC, HDC), and smooth muscle cells (SMCs; KRT18, TPM2), most prominent in Protocol II. Both protocols produced similar proportions of homeostatic MGL, while Protocol I gave rise to more proliferating/remodeling cells and Protocol II to more activated MGL and off-target cells. Differential expression analysis further distinguished the two protocols, with Protocol I cells enriched for genes linked to protein synthesis and cytokine/chemokine secretion, and Protocol II cells enriched for genes linked to intracellular transport, chronic stress response (TXNIP), and cytoskeletal dynamics (CYFIP1) (Fig. S1J).

To benchmark MGL against primary cells, we compared our data to a published human primary microglia scRNA-seq dataset ^46^ (Fig. S2). MGL from both protocols most closely resembled primary microglia from gestational week (GW) 9-15 (Fig. S2B), with Protocol II showing somewhat higher similarity to later time points and non-microglial lineages (Fig. S2F-H)(Fig. S2F-H). Despite expression of general myeloid markers, neither protocol yielded strong expression of the core microglial identity genes P2RY12, TMEM119, and SALL1. Given their overall similar transcriptomic profiles and the fewer off-target populations generated by Protocol I, we used it as the basis for subsequent MGL co-culture and optimisation.

### MGL co-culture with neural organoids increases similarity to primary microglia throughout the first month

Microglia identity is defined not only by its ontogeny, but also by its interaction with the brain environment ^47^. We hypothesized that co-culture of MGL with neural organoids would shift MGL phenotypes towards their in vivo counterparts. Moreover, analyzing this co-culture across multiple time points would provide a temporal view of how both MGL and the organoid co-evolve over time, revealing the dynamics and sequence of phenotypic changes as they unfold. Following Protocol I, we generated MGL-progenitors from a hiPSC line expressing green fluorescent protein (GFP) constitutively, in order to track MGL within the organoid and sort cells for subsequent experiments (Fig. S3A). Using the same parent cell line but without GFP expression, we produced unguided neural organoids, as described previously ^48^. On day 35 of organoid development, during neurogenesis, we cocultured organoids with MGL-precursors, with manual shaking and supplemented with FBS for the first 2 days to avoid MGL clumping (Fig. 1A, Fig. S3B). We supplemented the organoid media with IL-34 and GM-CSF for the first week to support unintegrated microglia during organoid infiltration. On day 3 of co-culture, many of the MGL had already attached to the organoids and by week 3, MGL integrated and acquired ramified morphology which was maintained for at least a further month of co-culture (day 58 of co-culture, final time point of the experiment) (Fig. 1B). MGL infiltrated the core of the organoid and were detected as IBA1-positively stained cells within the parenchyma of paraformaldehydefixed paraffin-embedded organoid slices (Fig. S3C).

**Figure 1.**
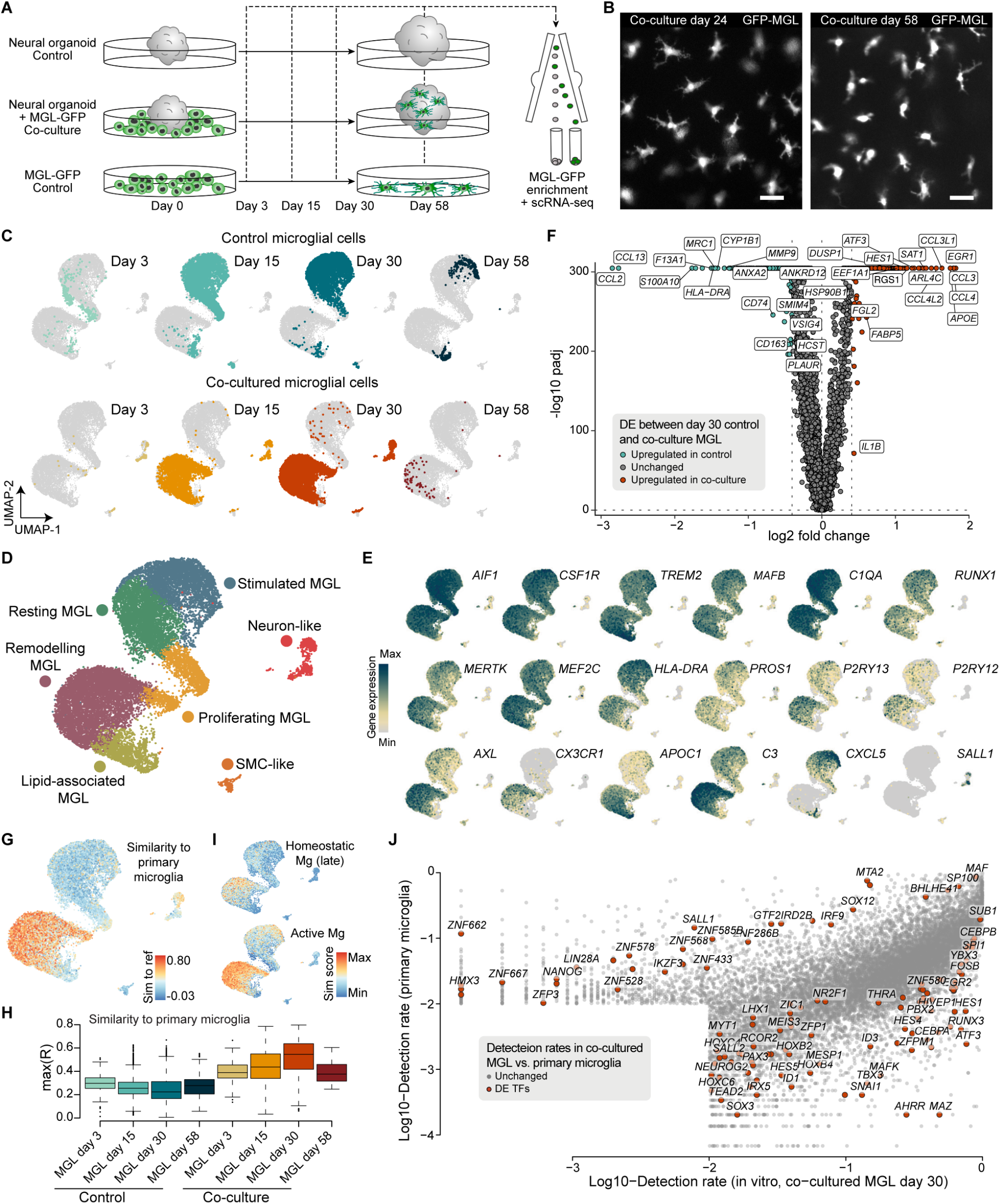
MGL co-culture within neural organoids increases similarity to human embryonic microglia. (A) Schematic representation of the co-culture setup. 35-day-old neural organoids were cultured separately (control), together with GFP-tagged MGL (GFP-MGL) precursors (co-culture) or GFP-MGL precursors were isolated and cultured as adherent cells in a dish (control). After 3, 15, 30 or 58 days, tissues or cells were enzymatically dissociated, FACS-sorted to enrich for GFP-positive cells and both GFP-negative and GFP-positive cells were subjected to scRNA-seq. (B) Confocal image of whole organoids (black) co-cultured with GFP-MGL (white) at day 24 and day 58 of co-culture. (C, D) UMAP representation of control (top) and co-cultured (bottom) MGL colored by time-point and co-culture (C), by cell type (D). (E) Feature plots of homeostatic, proliferative-region-associated microglia and border-associated macrophage marker genes in integrated co-cultured and control MGL. (F) Volcano plot of DE genes between day 30 of co-culture versus control MGL. (G, H) RSS similarity score of MGL to meta-cells of a published primary fetal human microglia scRNA-seq dataset 46 mapped onto the UMAP representation (G) or presented as a boxplot (H). (I) Module scores for late homeostatic or active primary microglia signature, based on highly expressed genes in primary fetal microglia ^46^, mapped onto UMAP of the MGL. (J) Scatter plot of (log10) detection rates of common genes in primary and 30-day co-cultured MGL, with DE TFs colored in red. Scale bar: B - 50 µm. MGL - induced microglia-like cells, DE - differentially expressed, SMC - smooth muscle cell, TF - transcription factor, Mg - microglia. See also figures S1 to S4.

On day 3, 15, 30 and 58 of co-culture, we dissociated control organoids, control MGL 2D mono-cultures and organoid-MGL co-cultures, enriched for sorted GFP-positive cells (MGL), and performed scRNA-seq (Fig. S3D). After filtering cells with low quality or potential doublets, we analyzed a total of 9,402 co-culture MGL, 10,161 control MGL, 25,119 co-culture organoid, and 15,890 control organoid cells (Fig. S3E). Normalized gene expression matrices were integrated using Cluster Similarity Spectrum (CSS 49), scaled and represented in 2D as a UMAP (Fig. 1C). MGL transcriptomic profiles exhibited a clear separation depending on the condition from which a given cell originated. Different subpopulations were further annotated based on unbiased clustering and DE gene expression (Fig. 1E, F, Fig. 2B, C, Fig. S4A). All MGL cells highly expressed typical microglia markers such as AIF1, CSF1R and TREM2 and both control and co-cultured cells contributed to a cluster of ‘proliferating MGL’.

**Figure 2.**
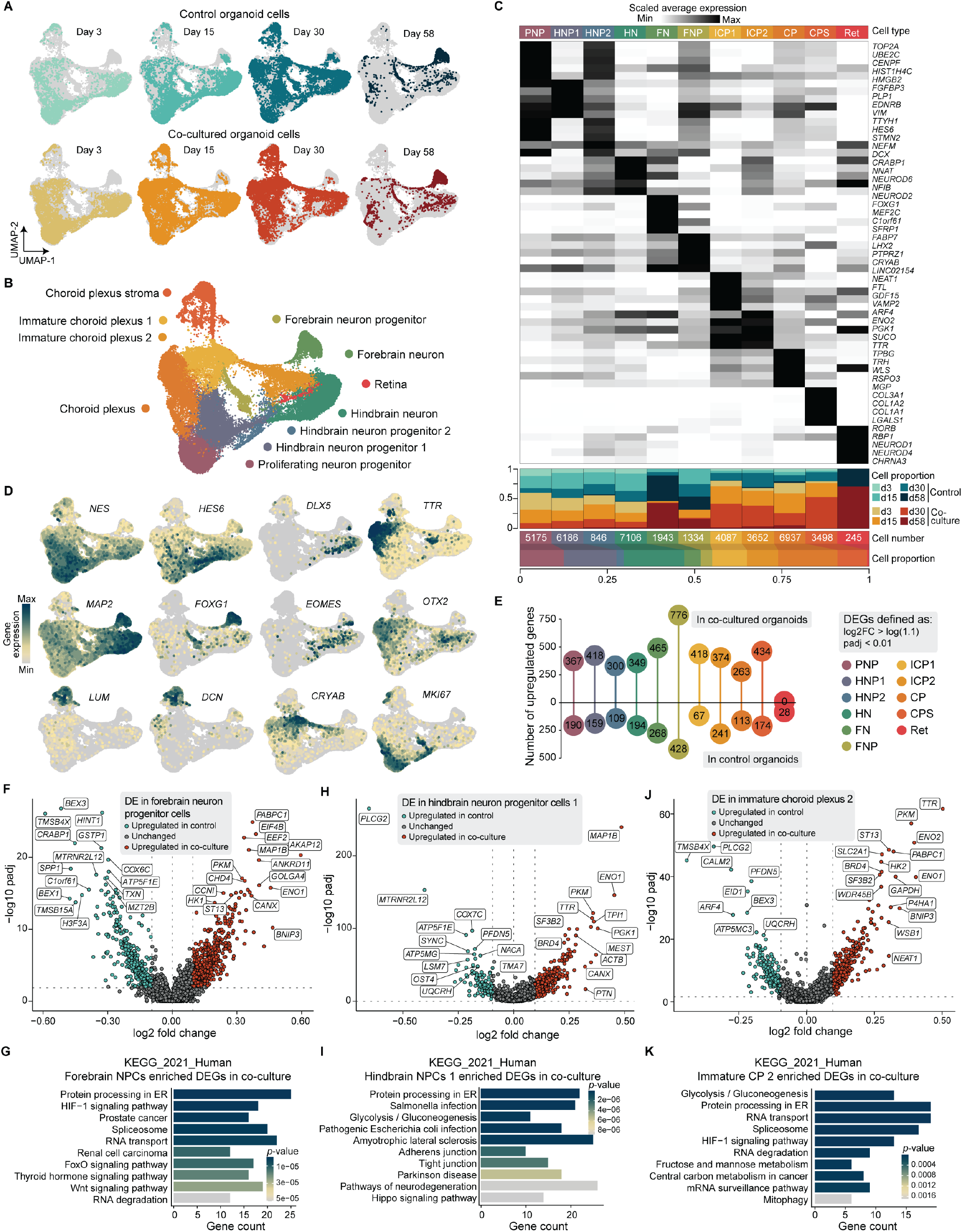
MGL cells influence neural cell states within brain organoids. (A, B) UMAP of integrated organoid transcriptomes cultured with or without MGL colored by sample origin (A) or by cell type (B). (C) Heatmap of scaled gene expression (z-score) of top 5 cell type markers combined with barplots showing the proportion of cells from each sample constituting each cell type and the total number and ratio of cells. (D) Feature plots of marker genes in organoids. (E) Lollipop plot showing number of DEGs in organoid cultured with versus without MGL across cell types. DEGs were defined as genes whose expression differs with log2FC > log(1.1) and have p-value adjusted < 0.01. (F, H, J) Volcano plot of DEGs between control and MGL co-cultured organoids in ‘Forebrain neuron progenitor cells’ (F), ‘Hindbrain neuron progenitor cells 1’ (H) or ‘Immature choroid plexus 2’ (J). (G, I, K) KEGG pathway enrichment analysis of processes mapping to genes upregulated in the co-cultured organoids in ‘Forebrain neuron progenitor cells’ (G), ‘Hindbrain neuron progenitor cells 1’ (I) or ‘Immature choroid plexus 2’ (K). DE(G) - differentially expressed (gene), d[…] - day […], PNP - proliferating neuron progenitor, HNP - hindbrain neuron progenitor, HN - hindbrain neuron, FN - forebrain neuron, FNP - forebrain neuron progenitor, ICP - immature choroid plexus, CP - choroid plexus, CPS - choroid plexus stroma, Ret - retina, KEGG - Kyoto Encyclopedia of Genes and Genomes.

Interestingly, co-cultured cells after 15 and 30 days largely formed clusters enriched in genes associated with homeostasis (P2RY13, CX3CR1) ^44^, synaptic pruning and clearing of apoptotic cells (C3, AXL) ^50,51^, which we called ‘remodeling MGL’, as well as in lipid-related genes (APOC1, APOE, ACP, increased TREM2), annotated as ‘lipid-associated MGL’ (Fig. 1E, F, Fig. S4A). Control MGL were marked by F13A1 (‘resting MGL’) and CXCL5 and HLA-DRA (‘activated MGL’) (Fig. S4A). Two small clusters segregated from the MGL populations. One, enriched in TPM2 and KRT18 was formed largely by day 15 control MGL and termed ‘smooth muscle cell (SMC)-like’ (Fig. 1E, Fig. S4A). The other one (‘neuron-like’) consisted of cocultured MGL and expressed typical neuronal markers such as TUBA1A and SOX4. As these cells also expressed, albeit at low levels, microglia markers, these MGL might represent a phagocytosing population, which have performed synaptic pruning and therefore engulfed neuronal RNA in the process. Neuronal gene-enriched microglia have been detected early during embryonic development ^44^, similar to the ’neuron-like’ subpopulation in our dataset.

DE analysis between control and 30 day co-cultured MGL (Fig. 1F, Table S3) showed that co-culture leads to upregulation of the chemokines CCL3, CCL4 and CCL4L2, which have previously been reported to be specifically associated with the human brain environment ^40,52^ and are expressed in primary microglia ^46^. Co-cultured MGL were also marked by increased expression of the early response genes EGR1, ATF3 and DUSP1, enriched in primary microglia ^15,46^. In the 2D MGL culture, there was increased expression of F13A1 and MRC1, which are expressed in BAMs and intermediate development stages of microglia ^40,53^, as well as the chemokines CCL2 and CCL13, S100A10 and ANXA2, previously implicated in cell migration ^54-57^.

Pseudobulk differential expression analysis (DESeq2 58) showed gradual increase in the expression of genes involved in Type I interferon pathway (IFI27, IFIT) over time in co-cultured microglia, whereas these factors were depleted in 2D cultured MGL (Fig. S4B, C) Non-microglia specific genes, such as CLDN1 or MYO10 exhibited reduced expression upon co-culture (Fig. S4B, C).

In order to compare MGL to human microglia, we used Reference Similarity Spectrum (RSS) ^49^ to calculate the similarity between MGL (query dataset) and to all microglia clusters from the human primary microglia (Mg) scRNA-seq dataset ^46^ (Table S4, genes used for RSS similarity calculations) introduced above (reference dataset) (Fig. S2) ^46^, excluding early microglia resembling monocytes, perivascular macrophages and SOX-high microglia cells (see Methods). Co-cultured MGL exhibited a transcriptional profile, substantially closer to that of Mg than the control cells (Fig. 1G, H), with strongest similarity between MGL and active primary developmental microglia at day 30 of co-culture (Fig. 1I, Fig. S4E-G). Furthermore, co-culture of 15 to 30 days resulted in an increase in MGL maturation as defined by their resemblance to Mg from embryos of different age (Fig. S4H, I).

An interesting observation was the decreased number of MGL recovered from brain organoids at day 58 of coculture, as well as their decreased similarity to primary microglia (Fig. 1C, H, Fig. S3E). Furthermore, SALL1 was not detected in any of the MGL, except for the ‘neuron-like’ and ‘SMC-like’ clusters (Fig. 1E). Reports that Sall1- deficient mice exhibit normal microglia colonization of the brain, but have a strong proinflammatory phenotype with detrimental consequences for neurogenesis and tissue homeostasis, hint towards an incomplete microglia differentiation which might be hampering co-culture efforts. Unlike the MGL cells, after standard pre-processing, integrated organoid cells from both control and co-cultured samples did not exhibit any major transcriptomic changes (Fig. 2A). Following Louvain clustering, based on DE genes, we annotated the resulting 11 subpopulations marked by the expression of forebrain-, hindbrain-, choroid plexus- and retina-specific genes (Fig. 2B-D, Tables S5-S7). We noted a large contribution of co-cultured organoid cells to the choroid plexus clusters (‘Immature choroid plexus 1’, ‘Immature choroid plexus 2’, ‘Choroid plexus’, ‘Choroid plexus stroma’) as well as to the ‘Retina’-like population (Fig. 2C).

Glycolysis-signature, protein processing in the endoplasmic reticulum and RNA transport and degradation ((Fig. 2F-K) had higher representation in co-cultured ‘Forebrain NPC’, ‘Hindbrain NPC 1’, and ‘Immature choroid plexus 2’, while genes upregulated in the control organoids were frequently associated with oxidative phosphorylation and pathways of neurodegeneration. MAP1B, a marker of neuron migration, axonal elongation and guidance, specifically in brain areas showing high synaptic plasticity, was significantly upregulated upon co-culture with MGL in forebrain and hindbrain NPCs (Fig. 2F, H). Another gene involved in brain development BRD4, which is expressed in response to neuron stimulation, was also enriched in some co-cultured organoid cells (Fig. 2H, J). On the other hand, genes upregulated in the control organoids were the regulators of nerve growth factor-dependent neuronal survival and differentiation BEX1 and BEX3, the phospholipase PLCG2, as well as PFDN5, a negative regulator of c-Myc, among others (Fig. 2F, H, J). Overall, the transcriptomic profiles of organoid cells cultured with or without MGL suggest that MGL might have a neuroprotective role in neural organoids. Despite the overall increased similarity to the primary microglia transcriptome, co-cultured MGL showed a reduced expression of some microglial genes, such as GREM1, SIGLEC8, and the TFs MTA2, IKZF3, SALL1 (Fig. 1E, Fig. S4D). Crucially, SALL1 was not detected in any of the MGL, except for the ‘neuron-like’ and ‘SMC-like’ clusters (Fig. 1E), and expression of the homeostatic marker P2RY12 remained low. This immature phenotype suggests that co-culture of MGL with neural organoids is not sufficient for MGL to transition from a CNS-associated macrophage state into an authentic in vivo-like microglia phenotype, indicating missing cues.

### An optimized 2D culture of MGL resembles closer co– cultured and primary microglia

MGL co-cultured with neural organoids exhibit increased similarity to primary developmental microglia, yet co-culture does not induce the expression of crucial microglia signatures such as SALL1. To evaluate MGL differentiation cues, we isolated MGL precursors and cultured them in control media (condition A) or control media supplemented with small molecules (conditions B-J) (Fig. 3A, B). We added TGF-β (homeostatic program) (B), CX3CL1 (maturation) (D), Activin A (a TGF-β family ligand) (E), CHIR99021 (Wnt activator) (F), CD200 (maturation) (H), cholesterol (survival) (I) or a combination of several cues (C, G, J). The conditions were chosen based on previous research on microglia and particularly on stem cell-derived microglia ^14,59-63^. After 16 days of culture, we enzymatically detached, labeled with distinct hashtag oligos (HTOs) 64, and pooled MGL for droplet-based scRNA-seq (10X Genomics). Demultiplexing based on HTO classification distinguished all conditions (Fig. S5A, B). After exclusion of cells with double or negative HTO classification and further quality control, a total of 15,195 cells were analyzed and pseudobulk principal component (PC) analysis confirmed low variability between biological replicates (Fig. S5C). Interestingly, despite clear morphological differences between the individual conditions (Fig. 3B), the treatments did not induce large shifts in the cell cycle status of MGL (Fig. S5D). Normalized data were scaled, clustered (Louvain), integrated using CCA, and represented as a UMAP (Fig. 3C). We identified 8 clusters (cluster 0-7) with expression profiles of conditions B and C congregating largely together, and condition J forming two nearly explicit clusters (Fig. 3C, Fig. S5F, G). All cells expressed highly the microglia markers AIF1, CSF1R and CLEC7A, but conditions including TGF-β (B, C, J) resulted in the strongest depletion of the macrophage markers F13A1 and SELENOP and led to enrichment of the complement component C3 (Fig. 3E, Fig. S5H, E). Importantly, MGL still lacked the expression of SALL1, regardless of condition (Fig. S5H).

**Figure 3.**
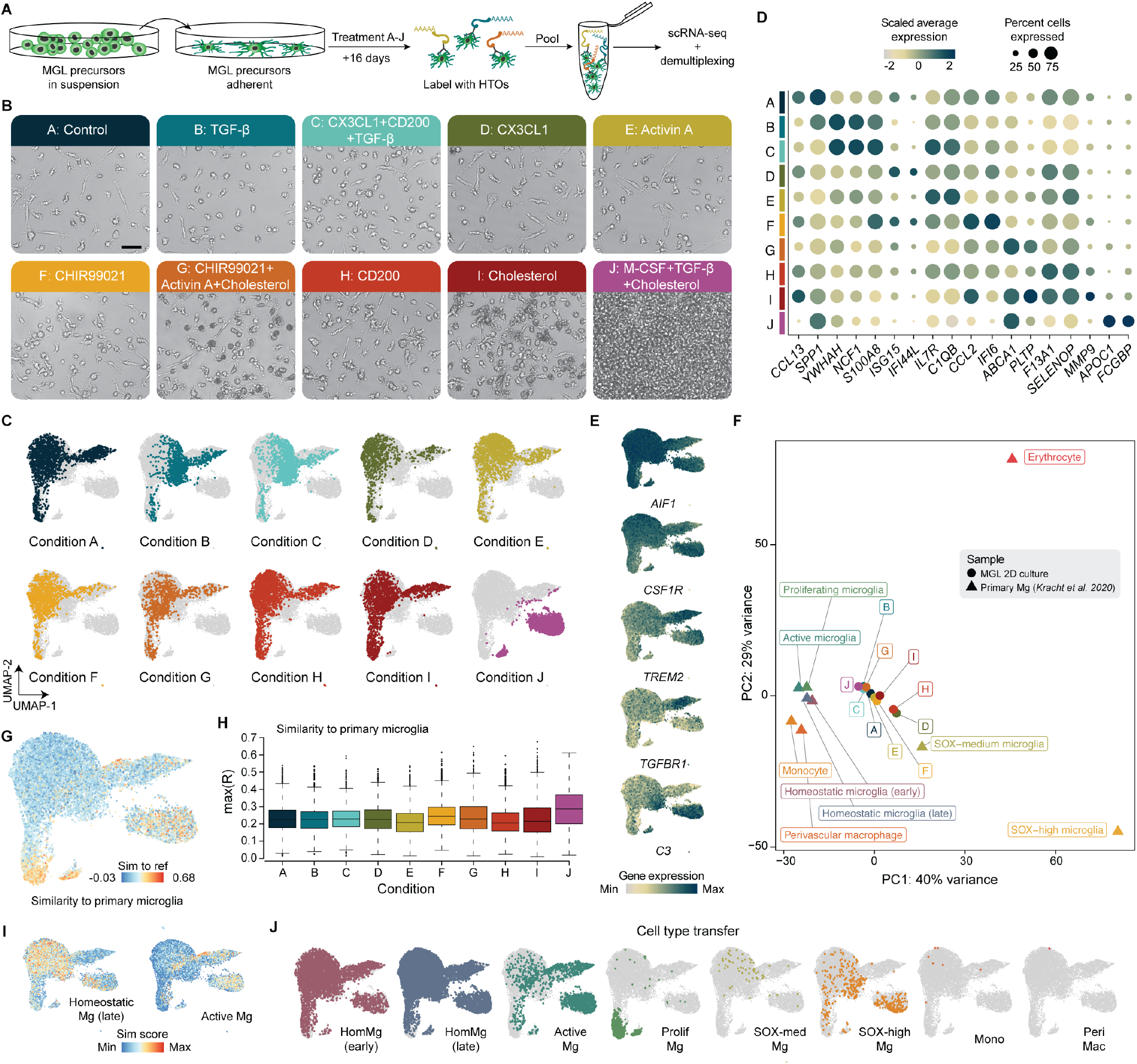
Growth factors direct MGL differentiation towards activated or homeostatic phenotypes. (A) Schematic representation of the experimental design of MGL culture optimization. MGL precursors were harvested, plated and treated with 9 different conditions for 16 days. Then, they were enzymatically dissociated, labeled with HTOs, pooled, subjected to scRNA-seq, and demultiplexed into individual conditions. (B) Brightfield images of MGL culture at day 16 and information about the small molecules added to the original protocol in each condition. (C) UMAP of MGL subjected to different treatments colored by condition. (D) Dotplot showing scaled gene expression (color) and percentage of cells expressing the respective gene (dot size) of top marker genes for each condition. (E) Feature plots of marker genes in MGL. (F) PC analysis plot of pseudobulk scRNA-seq data from MGL conditions or primary microglia cell subpopulations delineating transcriptomic similarity. (G, H) RSS similarity score of MGL to meta-cells of a published primary fetal human microglia scRNA-seq dataset 46 mapped onto the UMAP representation (G) or presented as a boxplot (H). (I) Module scores for late homeostatic or active primary microglia signature, based on highly expressed genes in primary fetal microglia 46, mapped onto UMAP of the MGL. (J) Label transfer of primary microglia 46 onto MGL. Scale bar: B - 50 µm. HTO - hashtag oligo, PC - principal component, Mg - microglia. See also figures S2 and S5.

Pseudobulk PC analysis of MGL conditions and human primary microglia ^46^ highlighted increased transcriptome similarity of TGF-β-treated MGL to primary developmental microglia subpopulations (Fig. 3F). These results were supported by RSS analysis of primary Mg meta niches 46 as the reference to MGL as the query dataset, with overall highest similarity score exhibited by MGL from condition J (Fig. 3G, H). Intriguingly, condition J [M-CSF, TGF-β, cholesterol] also contained the highest proportion of cells resembling microglia from older samples (GW18) (Fig. S5I-J) and activated microglia ((Fig. 3I, J, (Fig. S5K). Conditions B [TGF-β] and C [CX3CL1+CD200+TGF-β] enhanced the expression of homeostatic microglial markers of MGL (Fig. 3I, J, Fig. S5K).

In summary, these results uncovered conditions inducing homeostatic or activated microglia signatures with increased correlation to corresponding signatures observed in primary developmental microglia, in an in vitro setting. Still these effects were not sufficient to induce SALL1 expression or reach a similar resemblance to primary developmental microglia as was observed in MGL co-cultured within neural organoids for 30 days, suggesting that additional cues are still missing to fully recapitulate in vivo microglia phenotypes.

### SALL1 overexpression shifts MGL phenotype towards an early microglia state

To test whether SALL1 expression is the missing cue for generating more authentic microglia from pluripotency, we induced expression of SALL1 in MGL (Fig. 4A)). Based on a Tet-On system, we transfected cells with reverse tetracycline-controlled transactivator (rtTA) and with SALL1, fused to eGFP, under a tetracycline-responsive element (TRE) promoter (see Methods). This allowed us to induce SALL1 expression at a time point of choice by the addition of 2 µg/mL Doxycycline to the culture media. We generated double-positive (rtTA/SALL1) stem cells, differentiated them towards microglia and continuously induced SALL1 overexpression (OE) from day 8. On day 15 of microglia differentiation, we sorted GFP-positive (GFP+) MGL, and performed scRNA-seq (10X Genomics) of the selected cell population. As controls we used MGL, transfected with the Tet-ON system vectors but untreated with Doxycycline (Control noDox) and untransfected MGL treated with Doxycycline (Control noVector). After quality control, we analyzed 4,259 SALL1 OE cells, 5,694 control noVector cells and 5,605 noDox control cells. After normalizing, integrating, and scaling, Louvain clustering resulted in six major cell populations (Fig. 4B, Fig. S6B, C). Clusters marked by increased expression of BAM-associated genes, such as LYVE1 and F13A1, were annotated as “BAM_1” and “BAM_2”. Clusters enriched in SPP1 and GPNMB, among other PAM markers, were annotated “PAM_1” and “PAM_2”. Interestingly, cells in which SALL1 had been overexpressed showed reduced expression of cell cycle marker genes compared to controls, associated with a surveillant, quiescent phenotype. MGL in which SALL1 was overexpressed had an enriched PAM signature, exhibited reduced BAM signature and had variable expression of homeostatic markers (Fig. 4C). Differential gene expression analysis between control noVector and SALL1 OE MGL with DESeq2 58 revealed upregulation of activation markers such as MCEMP1, HLA-DRA, S100A9 and chemokines such as CCL3, CCL4, CXCL8 and CCL13 (Fig. S6F). Top GO terms for biological processes associated with these genes were “Antigen processing and presentation” and “Mitochondria respiratory chain complex” (Fig. 4D). The overall transcriptome similarity to primary microglia, calculated as Pearson correlation coefficient (see Methods) showed only a slight increase (Fig. S6E), which might also reflect the absence of homeostatic microglia expression signatures such as P2RY12 and TMEM119. Together, results from overexpression of SALL1 in 2D MGL monoculture suggest that although SALL1 is essential for microglia homeostasis, other cues are still missing.

**Figure 4.**
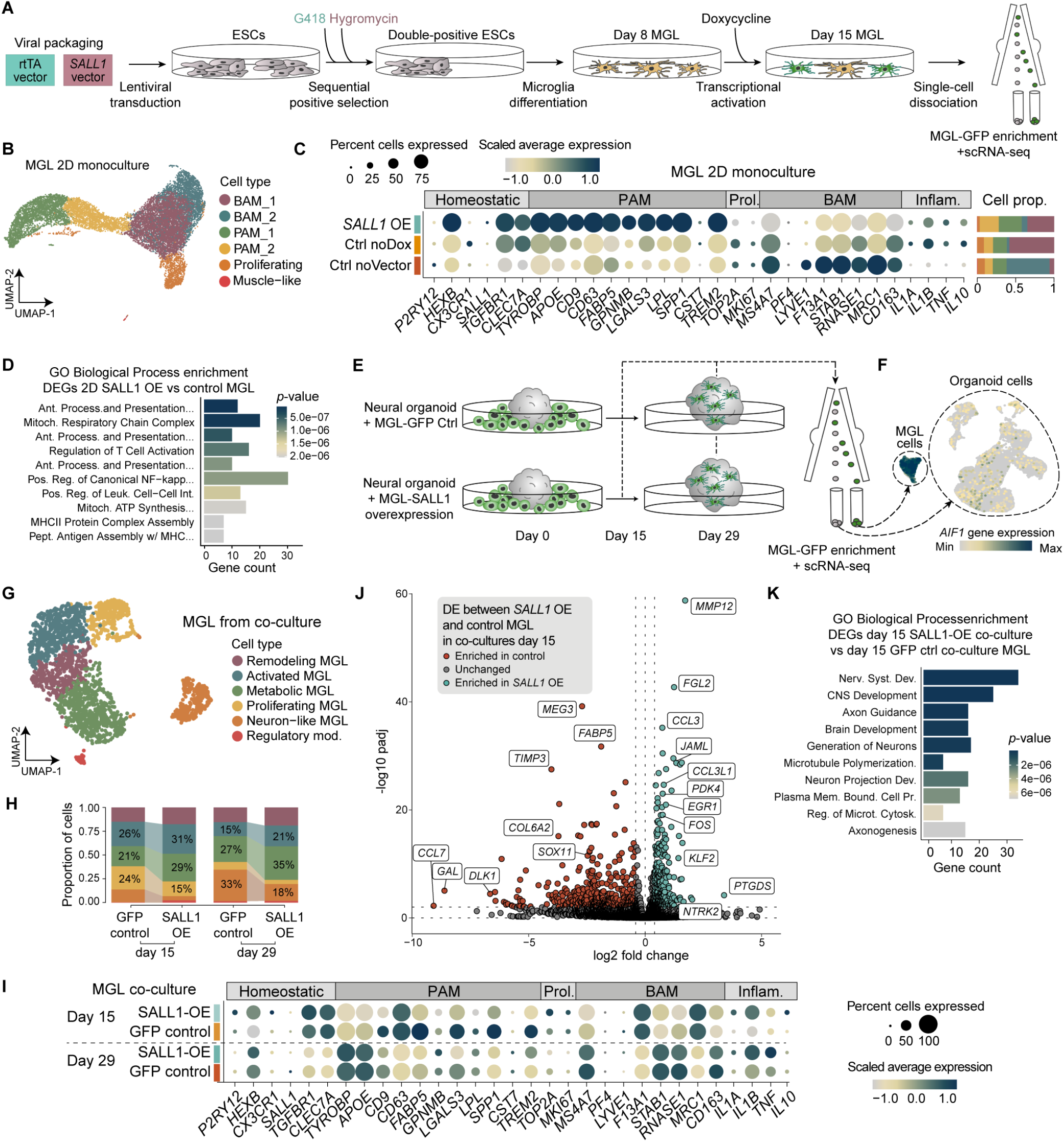
SALL1 overexpression in MGL induces transcriptional states resembling in vivo microglia. (A) Schematic representation of the experimental design of exogenous SALL1 overexpression in MGL using a Tet-On system. Two vectors were generated, packaged into lentiviral particles and used for ESC transfection. An rtTA vector, encoding a reverse Tet repressor and Neomycin resistance. SALL1 vector, coding for SALL1 fused to GFP, under the control of a TRE promoter and resistance to Hygromycin. Cells were transduced with lentivirus containing the plasmids and sequential selection of transfected cells was performed with G418 for rtTA, followed by Hygromycin for the SALL1 plasmid. Double-positive cells were differentiated into MGL. At day 8 of final microglia maturation, Doxycycline was added to the media for 7 days, thereby activating the rtTA system and leading to SALL1-GFP expression. Control MGL were not treated with Doxycycline (Control noDox) or not transfected with the vector but treated with Doxycycline (Control noVector). GFP cells were FACS sorted and control and GFP-positive cells were subjected to scRNA-seq. (B) UMAP of 2D monoculture MGL subjected to different treatments colored by cell type. (C) Dotplot showing scaled gene expression (color) and percentage of cells expressing the respective gene (dot size) of homeostatic, PAM, proliferating, BAM or inflammation-related markers for each condition in 2D monoculture. Stacked bar plots to the right show the proportion of each cell type for every condition. (D) GO analysis of genes upregulated in SALL1 OE MGL in comparison to Control noVector MGL in 2D monoculture. (E) Schematic representation of the experimental design of MGL-GFP or MGL-SALL1 OE GFP with neural organoids. (F) Feature plot of all organoid and MGL cells, sequenced with a droplet-based method, showing the expression of AIF1, a macrophage marker.. (G) UMAP of MGL (control or SALL1 OE, which have been co-cultured with neural organoids colored by cell type. (H) Barplot showing proportion of cells in each GFP control and SALL1 OE MGL co-cultured with organoids for 15 or 29 days, based on cell type. (I) Dotplot showing scaled gene expression (color) and percentage of cells expressing the respective gene (dot size) of homeostatic, PAM, proliferating, BAM or inflammation-related markers for GFP control and SALL1 OE MGL co-cultured with organoids for 15 or 29 days. (J) Volcano plot of differentially expressed genes between SALL1 OE and control MGL co-cultured with organoids for 15 days. (K) GO analysis of genes upregulated in SALL1 OE MGL in comparison to GFP control MGL in 15-day organoid co-culture. ESC - embryonic stem cell, rtTA - reverse tetracycline-controlled transactivator, GFP - enhanced green fluorescence protein, TRE - tetracycline-responsive element, scRNA-seq - single-cell RNA-sequencing, MGL - microglia-like cells, PAM - proliferative-region-associated microglia, BAM - border-associated macrophages, GO - gene ontology, DE - differential expression. See also figures S6 and S7.

### SALL1 overexpression in MGL co-cultured with neural organoids further shifts MGL phenotype towards a developing microglia state

SALL1 OE in MGL and MGL co-culture with neural organoids both increased phenotypic similarity to in vivo microglia. To test whether these effects are cumulative, we co-cultured day 31 or day 35 neural organoids with stem cell-derived microglia, overexpressing SALL1-GFP or only GFP as control, upon induction with Doxycycline. On day 15 and day 29 of co-culture, we dissociated co-cultures, FACS-sorted GFP+ and GFP-cells and performed scRNA-seq using two approaches - droplet-based 10X Genomics and plate-based MARS-seq.

After quality control, we analyzed 17,928 control and 16,705 SALL1 OE cells from the 10X data and 2,096 control and 2,966 SALL1 OE cells from the MARS-seq data. After normalization, integration using CSS, scaling and Louvain clustering, clusters enriched in AIF1-positive cells were subset and further analyzed as “MGL”, whereas all other clusters were further analyzed as “organoid” cells (see Methods) (Fig. 4F).

MGL subset data was scaled, integrated and clustered into six subpopulations, annotated based on highly expressed genes (Fig. 4G, Fig. S6G-I). SALL1 overexpression led to a decreased number of proliferating MGL and neuron-like cells in comparison to control, as well as to an increased number of metabolic and activated (expressing antigen-presenting cell markers) MGL (Fig. 4H). Similar effects on MGL fate decisions in SALL1 OE MGL were seen at both day 15 and day 29 of co-culture. SALL1 OE upregulated expression of some homeostatic markers (P2RY12, HEXB, CX3CR1) whereas the expression of many PAM genes was decreased in SALL1 OE MGL, e.g. CD9, FABP5, SPP1 (Fig. 4I).

SALL1 OE led to upregulation of genes associated with immune functions and inflammation, such as IL1A, IL1B, TNF, IL10 (Fig. 4I) and PTGDS, FGL2 (Fig. 4J) after 15 days of co-culture. DE analysis between GFP control and SALL1 OE MGL co-cultured with organoids for 15 days showed that co-culture leads to upregulation of the chemokines CCL3 and CCL3L1, which have previously been reported to be specifically associated with the human brain environment ^40,52^ and are expressed in normal microglia ^46^ (Table S8). Co-cultured MGL were also marked by increased expression of the early response genes EGR1, KLF2 and FOS, which prior studies have linked to a more vigilant or primed microglial state ^65^. GO terms associated with upregulated genes in SALL1 OE versus GFP control MGL were related to nervous system development (Fig. 4J, K, Fig. S6K) and caused time-dependent shifts in major signaling pathways (Fig. S6L). All four conditions were detected in all organoid cell clusters, upon integration and Louvain clustering (Fig. S7A-D). The “neural progenitor cell” cluster was of particular interest as we expected NPCs to be affected by communication with MGL. Among the DEGs between NPCs from SALL1 OE and control were genes with a neuroprotective role, such as MTRNR2L12, NTRK2, SLC1A3, APOE, CLU (Fig. S7E), and GO analysis associated many of the DEGs with protein processing (Fig. S7F). GO terms depleted in SALL1 condition were associated with OXPHOS and disease (Fig. S7F). GO terms derived from receptor-ligand interaction analysis corroborated the findings that SALL1 OE MGL co-cultured organoids were enriched in terms linked to neural development at day 15 (Fig. S7F).

Taken together, we present that SALL1 overexpression in MGL co-cultured with neural organoids shifted MGL fate from an early developing monocyte-like stage, towards a more accurate homeostatic-remodelling microglia state, better resembling an in vivo-like microglia.

## Discussion

Despite the successful development of several protocols for MGL differentiation, there is no consensus yet on what is the best approach for differentiating cells that recapitulate microglia phenotype in vivo and how these could be implemented in brain organoids ^63,66^. In this study, we uncover the heterogeneous cell subpopulations that MGL acquire in 2D monoculture. We steer MGL phenotype towards an in vivo-like developmental state via co-culture with neural organoids, and explore varying media compositions which guide MGL into a more accurate resemblance of active or more homeostatic cell states. We show that none of the tested conditions fully recapitulate primary developmental microglia and therefore exogenously overexpress the key microglia regulator SALL1 as a means of bridging that gap, which resulted in a shift from a general monocyte-like state towards a more accurate microglia fate.

We compared two previously published protocols for the generation of stem cell-derived microglia, which do not require any enrichment steps, such as FACS, and take a little over 3 weeks to differentiate into fully differentiated microglia-like cells. While cells from the two protocols exhibited similar morphologies, their propensity to develop into various microglia states or off-target populations differed, with a high percentage of cells from Protocol II becoming SMC-, neutrophil- or mast cell-like. Despite 2D MGL culture closely resembling the profile of primary microglia, some signature genes were scantily (P2RY12) or not at all (SALL1) detected, highlighting the importance of probing deeply individual cells.

Co-culture of MGL with neural organoids for 30 days led to a three-fold increase in their similarity to primary developmental microglia, when compared to 2D culture of the same duration. Among the upregulated genes were members of the apolipoprotein family (APOE, APOC1) and the complement gene family (C3), as well as immediate early genes (EGR1, RGS1), all of which have been detected throughout the human and mouse microglia developmental program ^49,46,67^. Markers of homeostatic microglia displayed higher expression in co-cultured MGL, suggesting that proteins embedded in the membrane of organoid cells or secreted by them are key to microglia maturation. Simultaneously, the number of cells resembling non-microglia myeloid cells, expressing genes associated with a microglia-BAM intermediate during mouse development ^53^, substantially decreased upon co-culture. Organoid cells exhibited increased expression of genes associated with neuroprotection and protein processing upon 30-day-co-culture with MGL. Long-term co-culture (over 30 days) resulted in loss of MGL cell number. These results corroborate previous findings that after an initial MGL cell number expansion, there is a gradual loss of microglia in the co-culture ^39^, which could be related to stress induced by the progressive necrosis in the organoid core ^68^.

Challenges in mimicking microglia in a dish stem from the peculiar embryonic origin of microglia that is distinct from other tissue macrophages, as well as their high sensitivity to their environment in vivo, essentially, shaping their functions. Thus, we hypothesized that an optimisation of the currently used MGL-derivation protocols by activating transcriptional programs more similar to that of primary microglia would result in an organoid-microglia co-culture that better represents the true brain environment. While most of the conditions tested induced slight shifts in MGL phenotype, the addition of TGF-β or a combination of it and other small molecules generated the strongest changes from the control condition, with the condition combining TGF-β with M-CSF and cholesterol leading to the largest shift in MGL phenotype towards primary microglia from all tested conditions. This could be explained with the importance of TGF-β for early microglia development and homeostasis and for maintenance of primary microglia ex vivo, which has been extensively described elsewhere ^15,25^. Furthermore, the addition of exogenous cholesterol has been shown as crucial for prolonged microglia survival ^13^. Overall, TGF-β caused upregulation of microglia signature genes such as C3, CX3CR1 and EGR1, but was not sufficient to induce SALL1 - a result fitting previous work showing that TGF-β had an overall positive effect on mouse microglia ex vivo, but no significant effect on SALL1 expression ^69^.

There is a growing body of evidence that SALL1 is a critical regulator of microglia identity, which gets upregulated in hematopoietic progenitor cells that enter the embryonic brain ^67,69^ and acts as both a transcriptional activator and repressor ^26^. Recent studies demonstrate that TGF-β directly activates a microglia-specific super-enhancer regulating Sall1 via SMADs, particularly SMAD4 ^15,26,69^. However, consistent with our findings, TGF-β alone is not sufficient to induce SALL1 expression - further aspects are the specific ontogeny of microglia (different from brain-associated macrophages ^69^) as well as additional unidentified brain environmental factors ^14,70^. Our approach was, therefore, to design a Doxycycline-inducible SALL1 expression system, which allows us to activate SALL1 transcription in MGL at a desired time point. MGL in which SALL1 had been overexpressed exhibited markedly reduced expression of cell cycle-related genes and of some myeloid markers, while a plethora of markers of pre-microglia/ early microglia (SPP1, APOE, TREM2) were upregulated. There was no significant increase in transcriptomic similarity to primary developmental microglia but a shift towards a more microglia and less nonmicroglia monocyte-like fate. Single-cell RNA-sequencing studies have identified embryonic microglial populations that co-express canonical microglial marker, such as P2ry12 and Fcrl1, alongside BAM-associated genes, such as Mrc1 (encoding CD206), Lyve1 and Ms4a7. These cells are predominantly located in regions like the embryonic cortical plate and intermediate zone during mid-gestation (around E15.5 in mice).

SALL1 OE in MGL co-cultured in a 3D environment with neural organoids induced a clear shift from a developing monocyte- or BAM-like phenotype towards a more mature homeostatic MGL, with downregulation of PAM and upregulation of remodeling / immune activation-linked markers. SALL1 shifts transcription but the exact effect depends on the environment and particular needs the cell has to meet at that time. SALL1 overexpression in microglia within neural organoids drives a transcriptional state characterized by heightened responsiveness, extracellular matrix remodeling, and signaling activity, which may be representing a more developmentally mature or functionally active microglial phenotype that supports neural circuit refinement.

In summary, in this study we capture the heterogeneity within 2D MGL monocultures derived from two different protocols. We show that MGL mature further upon coculture with neural organoids and resemble more closely primary developmental microglia transcriptomes after 30 days of co-culture, but lose this signature over long-term culture. Different culture conditions of MGL were able to induce gene expression programs closer to resembling those of more active or homeostatic primary microglia, or slightly increase overall transcriptomic similarity to developmental microglia. However, the largest transition towards a matured and more homeostatic microglial phenotype was achieved upon SALL1 OE in MGL in combination with co-culture with neural organoids, thus showing that SALL1 is an essential driver towards microglia fate and subsequent maturation. Our comprehensive studies will allow the community to choose their model system ranging from BAM-like to PAM-like microglia-like MGL, thus providing a novel toolbox for studying these subtypes in vitro.

### Limitations of the study

This work offers a wide and thorough transcriptomic exploration of pluripotent stem cell-derived microglia under different conditions. Important to note is that the genes induced in early microglia development overlap to a large extent with those upregulated in pathological conditions, making annotation of cell states challenging. Microglia and microglia-like cells exist in diverse and multidimensional states depending on the context and a unified nomenclature is lacking. The results add novel information to the pool of knowledge about the functional role of extrinsic (the environment, such as organoids) or intrinsic (gene expression of SALL1) factors in microglia cell specification and maturation. The milieu of the neural organoids provide a plethora of essential cues but are not able to recapitulate the human brain without vasculature, or other glial or immune cells. Thus, further studies are needed to explore the effect of SALL1 overexpression under more complex 3D conditions.

## Author Contributions

M. T. N., A. C., J. H., R. O. and J. K performed cell culture, designed and executed all 10X single-cell RNA-sequencing experiments. K.F., R.O. and M. T. N. performed MARS-seq experiments. P.W., J.R., K.F., M.T.N. performed immunostaining experiments and analysis. M. T. N. performed computational analysis with support from Z. H. for co-culture experiments and primary data analysis and support from K.F. for MARS-seq analysis. H. C. L. designed and provided original overexpression vectors. L.G., K.F. and I. A. helped design SALL1 experiments and interpret transcriptomic data. M. T. N., J. G. C., I. A., B. W. and B. T. designed the study. M. T. N. and B. T. wrote the manuscript with contributions from A.C., J.H., K.F., N.L., J.G.C., B.W., I.A.. All authors edited and approved the manuscript.

## Acknowledgements

We thank the Treutlein and Camp labs for helpful discussion. Particularly, Nadezhda Azbukina, who provided invaluable advice on transcriptomic data analysis. Illumina sequencing was done by Ina Nissen, Elodie Vogel Brucklen and Christian Beisel of the Genomics facility at D-BSSE, ETH Zurich. FACS sorting support was provided by Mariangela Di Tacchio, Aleksandra Gumienny, Renan Antonialli and Thomas Horn of the Single Cell Facility at D-BSSE, ETH Zurich. This study was supported by the Bavarian Ministry of Science and the Arts in the framework of the Bavarian Research Consortium ‘Interaction of Human Brain Cells’ (ForInter) network. Additional funding came from the DFG, German Research Foundation) 505539112; KFO 5024 (A01).

## Data and Code Availability

Raw sequencing data will be deposited into ArrayExpress. Processed data and the VCF files for demultiplexing will be deposited in Mendeley Data.

## Material and methods

No statistical methods were used to predetermine sample size. The experiments were not randomized. The investigators were not blinded to allocation during the experiments and outcome assessment.

## Experimental Materials and Methods

### Human pluripotent stem cell culture

In this study we used the human embryonic stem cell (hESC) line H9 (non-diseased female donor) obtained from WiCell 73, the human induced pluripotent stem cell (hiPSC) line 01F49i-N-B7 (short B7; nondiseased female donor), kindly provided by the lab of B. Roska (Institute of Molecular and Clinical Ophthalmology Basel) through a Material Transfer Agreement 74 and the hiPSC line WTC expressing mTagRFPT-CAAX obtained from the Allen Institute for Cell Science. The use of the hESC line H9 for generation of protocol II MGL was approved by the Robert Koch Institute (166. Approval according to the German Stem Cell Act – Investigation of the role of myeloid cells in inflammatory processes of the central nervous system in cell and tissue models derived from hES cells, to Beate Winner from 19.04.2021).

All cells and stem cell-derived cultures were incubated at 37°C and 5% CO2 in a humidified incubator. Stem cells were cultured in mTeSR Plus media (STEMCELL Technologies, 05825) supplemented with 50 U/mL Penicillin/Streptomycin (P/S) on tissue culture-treated plates pre-coated with hESC-qualified Corning Matrigel matrix (Corning, 354277) in KnockOut-DMEM/F12 (Gibco, 12660012). Cells tested negative for mycoplasma on a regular basis using the EZ-PCR Mycoplasma Detection Kit (Biological Industries, 20-700-20) and found to be negative. Media was exchanged every second day. Once per week, cells were passaged in small clumps with 0.5 mM EDTA (Invitrogen, AM9261) in Dulbecco’s phosphate buffered saline (DPBS) (Gibco, 14190250) for regular maintenance. After thawing, cells were passaged at least once before being used for microglia or organoid generation. On the first day of passaging on thawing of the cells, media was supplemented with 5 µM Rock inhibitor Y-27362 (STEMCELL Technologies, 72304).

### Human embryonic kidney cell culture

Human embryonic kidney (HEK) cells T293 were cultured in 10 cm tissue culture-treated Petri dishes in DMEM supplemented with 10% FBS (Sigma Aldrich, F2442), 50 U/mL P/S (Gibco, 15140-122) and 2 mM Glutamax (Gibco, 35050-061). Cells were passaged every week using TrypLE Express (Gibco, 12605-010).

### Microglia differentiation form hPSCs

Stem cell-derived microglia of Protocol I, used throughout this manuscript, were generated as described before 42. In brief, 2-3 days after passaging, PSCs were dissociated into single cells using TrypLE Express (Gibco, 12605-010). On day 0 of culture, 10,000 cells/well were seeded in a 96-well ultra low-attachment plate in 100 µL embryoid body (EB) media/well [mTeSR Plus media (STEMCELL Technologies, 05825) supplemented with 50 U/mL P/S (Gibco, 15140-122), 10 µM Y-27362 (STEMCELL Technologies, 72304), 50 ng/mL BMP-4 (Peprotech, 100-05ET), 20 ng/mL SCF (PeproTech, 300-07) and 50 ng/mL VEGF-121 (PeproTech, 100-20A)]. Plates were centrifuged for 3 min at 300x g and incubated at 37°C and 5% CO2 in a humidified incubator. 2 days later, half of the media was exchanged with fresh media. On day 4, EBs were transferred to a 6-well tissue culture treated plate (16 EBs/well) and cultured in 3 mL/well of primitive macrophage precursor (PMP) media [X-VIVO 15 (Lonza, BE02-060F), 2 mM Glutamax (Gibco, 35050-061), 50 U/mL P/S (Gibco, 15140-122), 55 µM β-mercaptoethanol (Gibco, 21985-023), 100 ng/mL M-CSF (PeproTech, 300-25), 25 ng/mL IL-3 (Cell Guidance Systems, GFH80-100)]. Media was exchanged fresh every 5-7 days. From day 14 on MGL precursors were harvested during media exchange. This was done by collecting the media, centrifuging it at 200x g for 5 min, resuspending the pelleted cells in RPMI 1640 media (Gibco, 11875-093) and plating at 180,000 cells/cm2 of a tissue culture-treated plate. After 1 h, MGL precursors had attached to the bottom and media was exchanged with complete MGL media consisting of RPMI 1640 (Gibco, 11875-093), 10Stem cell-derived microglia of Protocol II were differentiated into MGL via a hematopoietic progenitor cell (HPC) stage as previously described 43. For HPC generation, the STEMDiff hematopoietic kit (STEMCELL Technologies, 05310) was used according to the manufacturer’s protocol. PSCs were seeded as small clumps and cultivated in STEMDiff hematopoietic medium. HPCs were collected on day 12 of differentiation and resuspended in RPMI 1640 supplemented with 10% FBS (Gibco), 50 U/mL P/S (Gibco, 15140-122), and 10 ng/ml GM-CSF (PeproTech, 300-03). On day 14 of differentiation, HPCs were further seeded in maturation medium additionally containing 50 ng/ml IL-34 (PeproTech, 200-34) and differentiated for 2 weeks into MGL, with medium added every 3 days and one 50% medium change after 1 week. Analyses were performed on differentiated iMGL after 2 weeks.

### Generation of neural organoids

Neural organoids were generated as described before with minor modifications 75. In brief, 2-3 days after passaging, PSCs were dissociated into single cells using TrypLE Express (Gibco, 12605-010). On day 0 of culture, 3,000 cells/well were seeded in a 96-well ultra low-attachment plate in 150 µL/well of mTeSR Plus (STEMCELL Technologies, 05825), supplemented with 50 U/mL P/S (Gibco, 15140-122) and 50 µM Y-27362 (STEMCELL Technologies, 72304). On day 2, half of the media was exchanged fresh. On day 4, half of the media was exchanged fresh, omitting Y-27362. On day 6, once EBs had reached a size of approximately 500-600 µm in diameter and exhibited brightened surface tissue, indicating ectodermal differentiation, formation of primitive neuroepithelial was induced. This was done by transferring each EB in a 24-well ultra-low attachment plate and culturing it for 5 days in 500 µL/well of neural induction media [DMEM/F-12 (Gibco, 21330-038), 50 U/mL P/S (Gibco, 15140-122), 1X N2 supplement (Gibco, 17502-048), 1X Glutamax (Gibco, 35030- 061), 1X MEM-NEAA (Sigma Aldrich, M7145), 10 mg/mL Heparin solution (Sigma Aldrich, H3149)]. On day 11, EBs showed optically translucent surface tissue consistent with neuroectoderm. EBs were embedded in Matrigel (Corning, 356234) droplets, incubated at 37°C and 5% CO2 in a humidified incubator for 30 min to let Matrigel solidify and then transferred into a 6-well ultra-low attachment plate with 5 mL/well neural differentiation (nDiff) media without vitamin A (-VitA) [1:1 of DMEM/F-12 media (Gibco, 31330-038) and Neurobasal media (Gibco, 21103-049), supplemented with 50 U/mL P/S (Gibco, 15140-122), 0.5X N2 supplement (Gibco, 17502-048), 0.5X B27 supplement without vitamin A (Gibco, 12587-010), 1X Glutamax (Gibco, 35030-061), 55 µM β-mercaptoethanol (Gibco, 21985-023), 1:4000 human insulin solution (Sigma Aldrich, I9278), 0.5X MEM-NEAA (Sigma Aldrich, M7145). Media was exchanged fresh 2 days later. On day 15, organoids were transferred to a 10 cm Petri dishes (8 organoids/dish) in 9 mL/dish nDiff media with vitamin A (nDiff+vitA) [nDiff media where B27 contains vitamin A (Gibco, 17504-044)]. Dishes were incubated on an orbital shaker at 65 rpm with fresh media exchange every 3-4 days.

### Culture of neural organoids with or without MGL

For microglia-organoid co-culture experiments, 5-week old neural organoids were transferred to a 6-well ultra-low attachment plate (1 organoid/well) with 3 mL media/well. Co-culture media consisted of nDiff+VitA supplemented with 0.5% FBS for the first 2 days, 100 ng/mL IL-34 (PeproTech, 200-34) and 10 ng/mL GM-CSF (PeproTech, 300-03) for the first 7 days. All further culture was performed in nDiff+VitA only. PMPs were harvested, centrifuged at 200x g for 5 min and resuspended in 1 mL co-culture media supplemented with IL-34 and GM-CSF. 1,000,000 PMPs were added to each well with an organoid and incubated statically with manual shaking every 4-6 hours for the first 24 hours. After this, the co-cultured organoids were kept on an orbital shaker at 65 rpm. For control microglia culture, MGL precursors were harvested and cultured on a 6-well tissue culture treated plate in complete microglia media as described above. For control organoid culture, neural organoids were transferred to a 6-well ultra-low attachment plate and cultured in the same media and shaking/static conditions as the organoids to which MGL precursors had been added.

### Co-culture of neural organoids with SALL1 OE MGL

For SALL1 OE or GFP control MGL co-cultures with organoids, the same procedure as above was followed. Media changes between day 7 and day 15 of co-culture included 4 µg/mL Dox added to the media. Co-cultures were kept for 15 or 29 days until dissociation, FACS sorting and scRNA-seq experiment (10X and MARS-seq) were done.

### Immunohistochemistry and imaging

For immunohistochemistry, MGL were cultured on 12 mm diameter, 1 thickness glass coverslips (Neuvitro, GG-12-oz), pre-coated with 10 µg/mL poly-D-lysine (Sigma Aldrich, A-003-E). Cells were washed once with DPBS (Gibco, 14190250) and incubated for 10 min at room temperature in 4% w/v paraformaldehyde (PFA). After three 5-min washes with DPBS, coverslips were stained immediately or stored in PBS at 4°C. Blocking was performed for 1 hour at room temperature with blocking solution containing 5% normal donkey serum (Abcam, ab7475) and 0.3% v/v Triton-X 100 (Sigma Aldrich, T8787) in DPBS (Gibco, 14190250). Primary antibodies were added at respective dilutions in blocking buffer and incubated for 16-20 hours at 4°C. Cells were then washed three times for 5 min with DPBS and incubated for 1 hour at room temperature with secondary antibodies conjugated to fluorescent proteins diluted 1:1000 in blocking buffer containing 1:1000 4’,6-diamidino-2-phenylindole (DAPI, Thermo Fisher Scientific, 62248) for nuclei counterstaining, protected from light. After three 10-min washes with DPBS, coverslips were left to air-dry at room temperature. Dry coverslips were mounted onto SuperFrost Plus glass slides (Thermo Fisher Scientific, 1047002) with Mowiol (Sigma Aldrich, 81381) and left to air-dry before imaging.

Primary antibodies used: anti-CX3CR1 (rabbit, Abcam, ab8021) at 1:100 dilution, anti-CD45 (rabbit, Abcam, ab10558) at 1:1000, anti-IBA1 (goat, Abcam, ab5076) at 1:500.

Secondary antibodies used: AlexaFluor 488 anti-rabbit (donkey, Thermo Fisher Scientific, 406416), AlexaFluor 488 antigoat (donkey, Thermo Fisher Scientific, A11055).

### Image processing

Following image acquisition, each image underwent denoising using a non-local means algorithm. Gaussian noise levels were estimated for each image using the estimate_sigma() function from the skimage.restoration module. The denoising was implemented via the denoise_nl_means() function from the scikit-image package 76. The filtering strength parameter was set to 0.8 × σ, which also served as the standard deviation for noise reduction. The patch size and patch distance parameters were configured to 5 and 6 pixels, respectively. These parameters were selected to achieve effective noise reduction while preserving critical image features and structural details. For visualization purposes, images were intensity-scaled between the first and 99th percentiles to optimize contrast. Multi-image overlays were generated using the NumPy’s dstack() function 77 and rendered using Matplotlib 78.

Images were stitched with Fiji, Grid/Collection stitching, Grid: snake by rows, 10% overlap, and otherwise default parameters (Linear Blending, Regression threshold 0.3, Max/avg displacement threshold 2.5, Absolute displacement threshold 3.5, Compute overlap (otherwise use approximate grid coordinates).

To remove image artefacts from the overview images, rough masks were created to define the region of interest (ROI). The stitched DAPI images were first scaled between the first and 99th percentile, then smoothed using Gaussian filtering (σ = 100) via the filters.gaussian() function from the scikit-image library. Otsu thresholding 79 was applied to the blurred image using the threshold_otsu() function from the same package, yielding a binarized mask. Holes within the mask were filled using scipy.ndimage.binary_fill_holes() 80, and the largest connected object was selected to isolate the organoid slice. The selected region was subsequently dilated using scipy.ndimage.morphology.binary_dilation() to fully enclose the organoid. The mask then underwent a second round of Gaussian smoothing (σ = 50), followed by hole-filling as described above, defining the final ROI. For visualization purposes, pixel values outside the organoid mask were set to zero prior to image scaling.

### Single-cell dissociation of 2D microglia

For dissociation into single cells, microglia were washed with DPBS (Gibco, 14190250) and incubated with pre-warmed Accutase (Sigma Aldrich, A6964) for 3-5 min at 37°C. The reaction was stopped with Hanks’ balanced salt solution (HBSS) (Sigma Aldrich, 55021C), supplemented with 2% FBS (Sigma Aldrich, F2442). The cell suspension was centrifuged at 300x g for 5 min at 4°C, the cell pellet was resuspended in HBSS with 2% FBS, passed through a 40 µm Flowmi strainer, washed with HBSS with 2% FBS, centrifuged at 300x g for 5 min at 4°C, and resuspended in HBSS with 2% BSA. The last washing with HBSS with 2% FBS was repeated for a total of two washes. Cells were counted, spun down and either adjusted to appropriate concentration and loaded in a lane of a 10X Genomics microfluidic chip, or sorted based on a specific marker using fluorescence activated cell sorting (FACS), as described below.

### Single-cell dissociation of neural organoids and microglia-organoid co-cultures

Control organoids and organoid co-cultured with microglia were dissociated using the Neural Tissue Dissociation Kit P (Miltenyi Biotec, 130092628). The organoids were cut into four-six pieces with a scalpel, washed three times with HBSS (Sigma Aldrich, 55021C), and incubated in papain (enzyme mix 1) at 37°C for 15 min, followed by the addition of enzyme A (enzyme mix 2). Organoid pieces were triturated using wide-bore 1,000-mL tips and incubated for an additional approximately 30 min with triturations every 10 min. The reaction was stopped with HBSS with 2% FBS. The cell suspension was centrifuged at 300x g for 5 min at 4°C, the cell pellet was resuspended in HBSS with 2% FBS, passed through a 40 µm Flowmi strainer, washed with HBSS with 2% FBS, centrifuged at 300x g for 5 min at 4°C, and resuspended in HBSS with 2% FBS. The last washing with HBSS with 2% FBS was repeated for a total of two washes. Cells were counted, spun down and either adjusted to appropriate concentration and loaded in a lane of a 10X Genomics microfluidic chip, or sorted based on a specific marker using FACS, as described below.

### FACS of dissociated cell and tissue culture

All single-cell suspensions were kept in HBSS with 2% FBS at 4°C degree until and during FACS. Positive and negative enhanced green fluorescent protein (eGFP) cell populations were sorted from microglia control (positive eGFP control), organoid control (negative eGFP control) and organoid-microglia co-cultures. FACS experiments were performed on a BD FACS Aria III Cell Sorter using BD FACSDiva 8.0.1 Software with a 100 µm nozzle.

### Single-cell RNA-sequencing (scRNA-seq) experimental procedure

Microglia cells used for comparison of Brownjohn and McQuade (modified) protocols were processed using the Chromium 10X Genomics Single Cell 3’ v2 Reagent Kit (10X Genomics, 120237). For all other single-cell RNA-sequencing (scRNA-seq) experiments, the Chromium 10X Genomics Single Cell 3’ v3.1 Reagent Kit (10X Genomics, 1000268) was used. The respective reagents and microfluidic chips were used following 10X Genomics recommendations.

All single-cell suspensions were kept in HBSS with 2% BSA at 4 until loading on the 10X microfluidic chip. Cell suspensions were adjusted to optimal range (700-1,2000 cells/µL) whenever possible prior to loading on the chip. All subsequent steps were performed following the supplier recommendations. Quantification and quality control of complementary DNA (cDNA) libraries was performed using High Sensitivity DNA assays on an Agilent Bioanalyzer and a Fragment Analyzer. cDNA libraries from microglia cells used for comparison of Brownjohn and McQuade (modified) protocols were sequenced on a HiSeq 2500 (high output) platform in paired-end mode with 26/8/0/98 cycles. All other cDNA libraries were sequenced on a NovaSeq 6000 platform in paired-end mode with 29/8/0/91 cycles (Experimental overview of 10X experiments is in Table S9).

### Labeling with hashtag oligos (HTOs) followed by scRNA-seq

For scRNA-seq experiments of alternative culture media conditions, 2D MGL cultures were dissociated into single cells as described above. were dissociated. Then, all samples were stained using conjugated hashtag oligos, pooled, sequenced and demultiplexed with regard to their hashtag, based on the original protocol for cell hashing64. In brief, after filtering through a 40 µm Flowmi mesh, cells from each condition were suspended in Staining Buffer at a concentration of 7,777 cells/µl (final volume of 45 µl) to which 5 µl TruStain Fc Receptor Blocking Solution (Biolegend, 422301) was added. The cell suspension was incubated for 10 min at 4°C, followed by the addition of 2 µl (1 µg) TotalSeq anti-human Hashtag antibody (hashtag oligo, HTO) (individual for each sample; Biolegend, catalog numbers 394601, 394603, 394605, 394607, 394609, 394611, 394613, 394615, 394617, 394619, 394623) and incubation for 30 min at 4°C, with manual shaking of the tubes every 10 min. Cell suspensions were washed with 1 ml Staining buffer (0.5% BSA in DPBS) followed by centrifugation at 300 x g for 5 min. Washing steps were repeated for a total of three times. Finally, cells were counted, resuspended to 1,000 cells/µl and 10 µl of each sample were added to a final mix. Two lanes on the 10x microfluidic chip (Chromium Single Cell 3’ v3.1 kit) were loaded with 25 µl of the final mix, targeting a total of 25,000 per lane, and sequenced on a NovaSeq 6000 following the manufacturer’s recommendations.

cDNA libraries were prepared following 10X Genomics standard protocol, described in more detail above. For preparation of the additional hashtag libraries, the 80 µl supernatant from the cDNA cleanup was further size-selected with 70 µl SPRIselect reagent (2.0X) and eluted in a final volume of 40 µl Buffer EB. A PCR reaction was set up with 5 µl of the eluted hashtag fraction, 2.5 µl i5 dual index, 1.5 µl i7 dual index, 50 µl Kapa Hifi Hotstart Ready Mix (Kapa Biosystems, KK2602) and 40 µl nuclease-free water with the following program: 2 min at 98°C, 19 cycles of (20 sec at 98°C, 30 sec at 64°C, 20 sec at 72°C), 5 min at 72°C in a BioRad C1000 Thermal Cycler. The final PCR product was size-selected with 1.2X SPRIselect reagent, cleaned up and eluted in 40 µl water.

### MARS-seq

Individual cells were isolated using the Symphony S8 cell sorter (BD Biosciences) into 384-well plates preloaded with 100nl of lysis buffer, 3µl of mineral oil, and 20nM barcoded poly(T) primers for reverse transcription. After sorting, the plates were centrifuged, rapidly frozen on dry ice, and stored at –80°C. Single-cell RNA-seq libraries were generated using the SPID-seq protocol 81. In this approach, polyadenylated mRNA from each cell was reverse transcribed into cDNA with barcode incorporation, followed by amplification. The cDNA from each plate was then pooled, fragmented, and amplified to produce sequencing libraries compatible with the Illumina platform. Each resulting library was evaluated for DNA concentration and overall quality.

### Generation of SALL1-inducible MGL

#### Vector cloning

To prepare the transcription factor insert for the SALL1 overexpressing vector, SALL1 gene cDNA ORF clone (Genescript, Clone-ID OHu18577) was first linearized using the restriction enzyme PvuI-HF (New England Biolabs, R3150) and SALL1 was then amplified by PCR following the protocol of the Phusion High-Fidelity PCR Master Mix (Thermo Scientific, F531L). In short, nuclease-free water, 1X Phusion Master Mix, 0.5 mM forward primer (GGGGGTCGACATGTCGCGGAGGAAGCAAGC) and 0.5 mM reverse primer (GGGGTCTAGATTAACTCGTGACGATCTCCTTGCTG) were mixed and 3% DMSO (Sigma Aldrich, D2650-5X10ML) was added. The reaction was incubated according to this 2-step cycling protocol without annealing: Initial denaturation at 98°C for 30 sec, 30 cycles of denaturation at 98°C for 10 sec and extension at 72°C for 2 min, final extension at 72°C for 10 min.

The amplicon size was validated by a 1% E-Gel™ EX Agarose Gel (G402021) on an E-Gel Power Snap Electrophoresis System (Invitrogen) and the PCR reaction was cleaned up using the Macherey-Nagel NucleoSpin Gel and PCR Clean-up Kit (Thermo Fisher Scientific, 740609.250). After clean-up, amplified SALL1 and the eGFP-containing backbone of interest (unpublished) were each digested with the restriction enzymes SalI-HF (New England Biolabs, R3138) and XbaI (New England Biolabs, R0145) in 10x CutSmart buffer (New England Biolabs, B6004) at 37°C for 1 hour. The backbone was treated with Quick CIP (New England Biolabs, M0525) at 37°C for 1h to prevent re-ligation of the linearized plasmid DNA. Agarose gels were run for both products (0.5% gel for SALL1, 1% gel for the backbone), the bands of the correct size were cut out of the gel and extracted via the Macherey-Nagel NucleoSpin Gel and PCR Clean-up Kit. The cleaned backbone and cleaned SALL1 insert were ligated following the T4 DNA ligase protocol (New England Biolabs, M0202 and B0202). The vector and the insert were mixed in a 1:3 ratio in T4 DNA Ligase Buffer (10X) with T4 DNA ligase for sticky-end ligation and incubated at 16°C overnight.

Transformation of the ligation reaction was performed using NEB 5-alpha competent Escherichia coli (New England Biolabs, C2987U). 10% of ligation was added to a tube containing 100 mL (2 joined vials) competent cells and incubated on ice for 30 min. Afterwards the cells were heat shocked at 42°C for 45 sec and put back on ice for 2 min. 3x volumes of S.O.C medium (Invitrogen, 15544034) were added to the tube and incubated at 37°C in a bacterial incubator for 1 h. The transformed ligations were spread out onto 10 cm LB (Sigma Aldrich) petri dishes with 1:1000 Ampicillin (Sigma Aldrich, A9518-5G), in a 1:2:3 ratio using glass beads (Sigma Aldrich, 18406-500G). Appropriate controls containing a control ligation without backbone, a control ligation without insert and a control transformation with only competent cells were plated as well. The plates were incubated at 37°C in a bacterial incubator overnight.

To check for the right construct, colony PCR was performed for an appropriate number of colonies following the KAPA 2G Robust HotStart protocol (Kapa Biosystems, KK5005). Single colonies were picked with a P10 pipette and transferred in an individual PCR tube containing 10 mL nuclease-free water. Each colony was then resuspended by light stirring with the pipette tip, and the remaining cells on the tip were inoculated in a culture flask containing 3 mL 2X YT medium (Sigma Aldrich, Y2377-250G) with 1:1000 Ampicillin. To each PCR tube containing bacteria suspension was added a master mix of nucleasefree water, 1x KAPA 2G Buffer B, dNTP mix (0.2 mM each dNTP), 0.5 mM forward primer (GGGACAAAGTGGATGGACTACAAAGA and GGGCAACGAGATCTCCGTCATTCAG), 0.5 mM reverse primer (GGGCTTGTGGAGCAGAAGATCTGATAA and GGGCAGCGTATCCACATAGCGTAAA) and 0.5 units KAPA 2G Robust HotStart polymerase. The reactions were incubated on a cycler according to protocol (5 min initial denaturation at 95°C, 10 sec denaturation at 95°C, 10 sec annealing at 60°C, 10 sec extension at 72°C and 30 cycles). As PCR readout 1 mL of each product was run on a 2% E-Gel™ EX Agarose Gel on an E-Gel Power Snap Electrophoresis System and only the clones showing the right band size were further incubated on a bacterial shaker at 37°C overnight. The DNA of the successful transformants was isolated according to the manual of the QIAprep Spin Miniprep Kit (Qiagen) and eluted in 50 mL elution buffer (10 mM Tris·Cl, pH 8.5). The concentration was measured via Nanodrop and the construct was checked by control restriction digestion with SalI-HF and XbaI before the DNA was sent to Microsynth (www.Microsynth.com) for Sanger sequencing.

### Lentivirus packaging

For lentivirus production, HEK293T cells were transfected with TransIT-293 Transfection Reagent (Mirus Bio, MIR2705). Before transfection, cells were passaged at a 1:2 ratio for at least 1 passage to ensure a good condition and seeded to achieve 80-90% confluency the next day. On the afternoon of day 1, the standard medium of the cells (DMEM [Gibco, 10569-010] + 1% GlutaMAX [Gibco, 35050-061] + 1% Pen/Strep [Gibco, 354277] + 10% FBS [Sigma Aldrich, F2442]) was exchanged for 10 mL fresh DMEM with low Pen/Strep concentration (DMEM + 1% GlutaMAX + 0.1% Pen/Strep + 10% FBS). 2 hours after medium change, the transfection reaction was assembled, consisting of the appropriate amounts of reverse tetracycline-controlled transactivator (rtTA) construct (pLVX-EF1a-tetOn-IRES-G418, Addgene, 84776), the eGFP-containing control backbone (unpublished) or the eGFP-SALL1 construct (see above), envelope plasmid (VSVG), packaging plasmids (Gag/Pol, Rev, Tat) and TransIT-293 Transfection Reagent in 1 mL Opti-MEM (Gibco, 31985062). After assembly, the mix was incubated at room temperature for 17 min and then added dropwise to the cells. The dish was gently shaken in a figure-8 motion to ensure proper mixing. On the morning the next day, media was exchanged for 6.5 mL of high FBS DMEM (DMEM + 1% GlutaMAX + 0.1% Pen/Strep + 30% FBS). On day 3 in the afternoon, the virus was harvested by collecting the suspension and spinning it down at 2000 rpm for 5 min. To remove cell debris, the clarified supernatant was transferred to a new tube. For virus concentration, 3 volumes of supernatant were combined with 1 volume of Lenti-X concentrator (Takara Bio, 631232) and the mixture was incubated at 4°C overnight. After 45 min of centrifugation at 1500 x g at 4°C, the supernatant was carefully removed from the off-white pellet. The pellet was resuspended 1:50 the original volume in PBS. The virus was stored at -80°C in single-use aliquots.

### Generation of a SALL1-inducible ESC line

To establish a stable rtTA/SALL1 cell line, H9 ESCs at 70%-80% confluency were split using TrypLE (Gibco, 12605-010) and seeded at 300,000 cells/well into a 6-well Matrigel-pre-coated plate (Corning, 354277) with mTeSR Plus media (STEMCELL Technologies, 100-0276), supplemented with CloneR2 (1:10) and 50 U/mL P/S (Gibco, 15140-122). No virus (control wells) or 10 µl, 20 µl or 40 µl of the rtTA-virus and of the SALL1-virus each were added per well. On the next day, media was exchanged with fresh mTeSR Plus with 50 U/mL P/S. On the next day, cells (70%-90% confluent) were split 1:20 using TrypLE into 10 cm Matrigel-pre-coated cell culture dishes in mTeSR Plus with 5 µM Y-27632 (STEMCELL Technologies, 72304). On the next day, positive selection for rtTA-positive cells was started by adding G418 disulfate salt (250 µg/mL final concentration, Sigma Aldrich, A1720-1G) to the culture media. After 4 days all control cells (wild type non-transfected cells with G418) were dead. Surviving transfected cells were split, as described above, and selection for SALL1-vector-positive cells was started by adding Hygromycin (100 µg/mL final concentration, Gibco, 10687010) to the culture media. After all control cells were dead, double-positive selected cells were split as described above or cryopreserved in mFreSR (STEMCELL Technologies, 05854).

### Culture of SALL1 OE MGL

MGL generation was performed as described above. 8 days after plating MGL in complete microglia media, Doxycycline (4 µg/mL final concentration, Sigma Aldrich, D9891-1G) was added to the media and kept until day 15 after plating, when eGFP-positive transfected cells and negative control cells were FACS sorted and scRNA-seq was performed, as described above. The same procedure was performed for eGFP-only control MGL.

## Quantification and Statistical Analysis

### Preprocessing and analysis of public primary fetal microglia dataset

Single-cell RNA-sequencing data of primary fetal microglia used as a reference in this article was previously published by Kracht et al.46 and retrieved from NCBI GEO, accession number GSE141862. The scRNA-seq data was then converted into a Seurat object (v4.0.0)82 and additional quality control was done by filtering cells with three times above or below the mean absolute deviation of the log10 transcript number detected or with mitochondrial transcript percentage higher than 10%. Data normalization was performed, the top 5,000 highly variable genes were defined using the vst method and mitochondria and ribosome genes were excluded from those. Data scaling was performed regressing out number of transcripts detected, percentage of mitochondria genes and percentage of ribosome genes. Principal component analysis (PCA) was applied to the scaled expression matrix for the first 50 principal components (PCs). Cells were clustered using Louvain clustering (resolution = 0.5). The cluster markers were identified using the Seurat package FindAllMarkers using Wilcoxon Rank Sum test requiring minimum 25% of cells in the cluster to express the gene with log2 fold change > log(1.2). Clusters were annotated based on canonical gene marker expression.

Only cells annotated as clearly microglia (“Homeostatic microglia (early)”, “Homeostatic microglia (late)”, “Activated microglia”, “Proliferating microglia”, “SOX-medium microglia”) were subset for generation of a microglia-similarity score and for differential detection rate calculation. The data of the subset cells were normalized, top 3,000 variable features were identified using vst method, scaled, Louvain clustering (resolution = 10) was performed and PCA was applied. For the generation of high-resolution clustering, dimensionality reduction embedding with Uniform Manifold Approximation and Projection (UMAP) was generated using the uwot (v0.1.14)83 package with “cosine” s distance metric for finding nearest neighbors, with local neighborhood size 30L, number of dimensions 2L, learning rate 1, local connectivity 1L, repulsion strength 1, and negative sample rate 5. Average gene expression of high resolution clusters was calculated and added onto the previously generated UMAP model.

### Microglia protocols comparison data preprocessing and analysis

Cell Ranger (10x Genomics, v4.0.0) was used to map the sequencing reads to the human reference (GRCh38-based, 10x Genomics, v3.0.0) and call cells to generate the count matrices. Next, the scRNA-seq data from MGL generated by the two different protocols was merged and normalized using Seurat (v4.0.0)82. Additional quality control was done by filtering cells with more than 9,000 genes detected or with mitochondrial transcript percentage higher than 15%. The top 3,000 highly variable genes were defined using the vst method. The two datasets were integrated using Canonical Correlation Analysis (CCA)84, part of the Seurat package, using the first 20 dimensions in the anchor weighting procedure. The expression values of each highly variable gene across cells were scaled, after regressing out the cell-cycle scores (G2M and S) calculated by Seurat, as well as the number of genes detected, the number of transcripts detected and the percentage of mitochondria genes. PCA was applied to the scaled expression matrix for the first 30 PCs. The first 20 dimensions were then used to generate the Uniform Manifold Approximation and Projection (UMAP) embedding, as well as clusters using Louvain clustering (resolution = 0.2). Cluster markers were identified using the Wilcoxon test as part of the Seurat function FindAllMarkers, filtering for genes expressed in at least 25DE and enrichment analysis between MGL subpopulations and all other MGL cell populations was performed using the DEenrichRPlot function provided by Seurat. The Wilcoxon Rank Sum test was used for identifying DEGs with p-value < 0.05 and log2 fold change > 0.3 using the Reactome 2022 pathway enrichR database85. To find DEGs between MGL generated with the two protocols, cells annotated as “smooth muscle cell”, “neutrophil” and “mast cell” were excluded and pseudobulk samples were generated across clusters for each sample by aggregating the row counts. Differential expression was calculated using the R package DESeq258, with design = celltype + protocol.

Seurat label transfer was used to transfer categorical information about microglia age in weeks and cell subtype by projecting the reference microglia dataset onto the query data (in vitro generated MGL). Anchors between the reference and query data were found with FindTransferAnchors via reciprocal PCA (rPCA) using the top 50 PCs on the reference and with the top 50 dimensions from the reduction to specify the neighbors search space. Label transfer based on timepoint or cell subtype was performed with TransferData using the internal PCA on the query only and the top 50 dimensions for the in the anchor weighting procedure.

### MGL co-culture data preprocessing and analysis

Control and co-culture MGL scRNA-sequencing reads were mapped to the human reference (GRCh38-based, 10x Genomics, v3.0.0) and cells were called to generate the count matrices using Cell Ranger (10x Genomics, v4.0.0). Samples from day 58 co-culture and control MGL and organoid had been loaded together on the same 10X Genomics lane and demultiplexed based on single nucleotide polymorphisms, as they belonged to two different cell lines - hiPSC B7 for MGL and hiPSC WTC for organoid cells, using demuxlet 86.

Data was further analysed using Seurat (v4.0.0). Quality control was applied by excluding cells with > 20% of mitochondria genes detected and further subsetting based on different parameters for each sample due to large sequencing depth variety between samples:

- Control MGL day 3: number of genes > 2,000 & < 9,000 & number of transcripts < 90,000;
- Control MGL day 15: number of genes > 1,500 & < 8,000 & number of transcripts < 55,000;
- Control MGL day 30: number of genes > 1,000 & < 7,000 & number of transcripts < 50,000;
- Control MGL day 58: number of genes > 2,000 & < 7,000 & number of transcripts < 50,000;
- Co-culture MGL day 3: number of genes > 2,000 & < 8,500 & number of transcripts < 100,000;
- Co-culture MGL day 15: number of genes > 1,500 & < 7,000 & number of transcripts < 50,000;
- Co-culture MGL day 30: number of genes > 1,000 & < 7,000 & number of transcripts < 50,000;
- Co-culture MGL day 58: number of genes > 1,000 & < 7,000 & number of transcripts < 50,000.

The scRNA-seq data of all MGL samples was merged, followed by normalization, highly variable gene identification (vst method, 3,000, mitochondrial genes and ribosomal genes excluded), data scaling (cell-cycle scores, percentage of mitochondria genes and number of detected genes regressed out) and PCA. CSS was further applied to integrate data of different samples, followed by PCA on CSS to obtain the PCA-reduced CSS representation (with 30 dimensions), which was then used to generate the UMAP embedding as well as Louvain clustering (resolution = 0.3). Cluster markers were calculated using FindAllMarkers function from Seurat with a Wilcoxon test, thresholding for expression in minimum 25% and logFC > 1.2. The resulting clusters were annotated based on the combinatorial expression of canonical cell type markers.

Differential gene expression between control and co-culture MGL at day 30 was calculated with Wilcoxon Rank Sum test from the presto package 87 (v1.0.0) and genes with absolute logFC > 1.5 and Benjamini-Hochberg (BH)-adjusted p-value < 0.01 were differentially expressed between the two conditions.

Differential detection rates between co-cultured MGL at day 30 and primary human microglia were calculated using binomial group test. First, smooth muscle cell-like and neuron-like cells were excluded from the MGL Seurat object. Only genes detected in both the primary and the in vitro MGL dataset were used for further analysis. Then, similarity values between primary and in vitro microglia were binarized and binomial regression was used to compare the frequency of both, including the cell types as a covariate. Mitochondria and ribosomal genes were excluded from the final differential detection results and only genes with BH-adjusted p-value < 0.05 were included.

Label transfer onto MGL based on cell type or age of primary microglia was performed using the Seurat package. First, with the function FindTransferAnchors, a set of anchors between the reference primary and the query in vitro datasets was generated, with reciprocal PCA, computing 50 PCs on the reference and using the first 50 dimensions from the reduction to specify the neighbor search space. Then, with the function TransferData, label transfer was performed onto the query dataset using the query PCA only by constructing a weights matrix defining the relationship between every query cell and every anchor, and multiplying it by a binary classification matrix, where each row corresponds to a possible class and each column corresponds to a reference anchor.

Similarity scores of control and co-cultured MGL to primary microglia were generated by calculating the Reference Similarity Spectrum (RSS) (simspec package 49 v0.0.0.9000) between the normalized in vitro MGL gene expression and pseudobulk normalized gene expression of the primary microglia with Spearman’s rank correlation and otherwise default parameters. For differential gene expression analysis based on the negative binomial distribution between control and co-culture MGL as well as between control samples at different time points, pseudobulk samples were generated and analysed using the package DESeq2 58 (v1.38.3). The full model included condition, timepoint and the interaction between timepoint and condition, while the reduced model included timepoint and condition only. The likelihood ratio test was then performed on the difference in deviance between the two models. Genes with abs(logFC) > 1.5 and BH-adjusted p-value < 0.05 (Wald test) were considered differentially expressed.

### Organoid co-culture data preprocessing and analysis

Control and co-culture MGL scRNA-sequencing reads were mapped to the human reference (GRCh38-based, 10x Genomics, v3.0.0) and cells were called to generate the count matrices using Cell Ranger (10x Genomics, v4.0.0). Samples from day 58 co-culture were processed with demuxlet as described above.

Data was further analysed using Seurat (v4.0.0). Quality control was applied by subsetting based on different parameters for each sample due to the variety between samples:

- Control organoid day 3: number of genes > 500 & < 7,500 & number of transcripts < 50,000 & percent.mt < 15;
- Control organoid day 15: number of genes > 1,000 & < 6,000 & number of transcripts < 35,000 & percent.mt < 5;
- Control organoid day 30: number of genes > 1,000 & < 8,000 & number of transcripts < 45,000 & percent.mt < 12;
- Control organoid day 58: number of genes > 1,800 & < 5,000 & number of transcripts < 20,000 & percent.mt < 15;
- Co-culture organoid day 3: number of genes > 500 & < 6,500 & number of transcripts < 35,000 & percent.mt < 12;
- Co-culture organoid day 15: number of genes > 1,000 & < 7,500 & number of transcripts < 35,000 & percent.mt < 12;
- Co-culture organoid day 30: number of genes > 500 & < 6,000 & number of transcripts < 45,000 & percent.mt < 12;
- Co-culture organoid day 58: number of genes > 1,800 & < 5,000 & number of transcripts < 22,000 & percent.mt < 15.

The scRNA-seq data of all organoid samples was merged, followed by normalization, highly variable gene identification (vst method, 3,000, mitochondrial genes and ribosomal genes excluded), data scaling (cell-cycle scores, percentage of mitochondria genes and number of detected genes regressed out) and PCA. CSS was further applied to integrate data of different samples, followed by PCA on CSS to obtain the PCA-reduced CSS representation (with 30 dimensions), which was then used to generate the UMAP embedding as well as Louvain clustering (resolution = 0.2). Cluster markers were calculated using FindAllMarkers function from Seurat with a Wilcoxon test, thresholding for expression in minimum 25% and logFC > 1.2. The resulting clusters were annotated based on the combinatorial expression of canonical cell type markers.

Differential gene expression between control and co-culture organoid in different clusters was calculated with Wilcoxon Rank Sum test from the presto package 87 (v1.0.0) and genes with absolute logFC > 1.1 and BH-adjusted p-value < 0.01 were differentially expressed between the two conditions. Pathway enrichment analysis of DEGs between control and co-culture in various cell types was performed with the R package enrichR 85 (v3.4) with default settings and extracting information from the KEGG 2021 database.

### Microglia protocol optimization data preprocessing and analysis

In order to demultiplex the cells based on their condition of origin, first HTO matrices were generated using CITE-seq-Count (v1.4.5)88. CITE-seq-Count outputs the number of unique molecular identifiers and read counts mapping to an antibody-oligo conjugate based on the sequencing data. Then, using Seurat, the HTO information was added to the Seurat object as an independent assay and used for demultiplexing with the function HTODemux with default parameters. Cells classified as singlets were assigned to the HTO of origin with the highest signal detected, while doublets and ambiguous cells were excluded from further analysis.

Cell Ranger (10x Genomics, v4.0.0) was used to generate transcriptomic count matrices of the MGL cultured under alternative culture conditions. Cells with mitochondria gene percentage higher than 20%, fewer than 600 detected genes and fewer than 1,000 detected transcripts were filtered out of the data. The scRNA-seq data of these samples were then merged and the cell cycle score was calculated with the Seurat function CellCycleScoring, followed by normalization, highly variable gene identification (vst method, 3,000), data scaling (cell cycle score regressed out) and PCA to obtain 50 PCs. CSS was further applied to integrate data of different samples, followed by PCA on CSS to obtain the PCA-reduced CSS representation (with 30 dimensions), which was then used to generate the UMAP embedding as well as Louvain clustering (resolution = 0.3). Cluster markers were calculated using FindAllMarkers function from Seurat with a Wilcoxon test, thresholding for expression in minimum 25% and logFC > 1.2.

Similarity scores of control and co-cultured MGL to primary microglia were generated by calculating the RSS (simspec package 49), as well as label transfer from primary microglia to MGL were generated as for the co-cultured MGL (see above). For differential gene expression analysis based on the negative binomial distribution between different culture conditions at different time points, pseudobulk samples were generated and analysed using the package DESeq2 58 (v1.38.3) with design = replicate + condition and genes with abs(logFC) > 1.5 and BH-adjusted p-value < 0.01 considered as DE.

### SALL1 OE MGL monoculture data preprocessing and analysis

Cell Ranger (10x Genomics, v4.0.0) was used to map the sequencing reads to the human reference (GRCh38-based, 10x Genomics, v3.0.0) and call cells to generate the count matrices. Next, the scRNA-seq data from MGL generated by the two different protocols was merged and normalized using Seurat (v4.0.0)82. Additional quality control was done by filtering out cells with less than 1,000 genes detected or with mitochondrial transcript percentage higher than 30%. Mapping of transcripts to the gene for Hygromycin resistance was used to infer vector expression within cells. As Hygromycin-resistance gene was also detected in the transcripts from control cells which had been transfected with the vector but its expression had not been induced with Dox, we suspected potential leakage. We therefore excluded control cells in which Hygromycin-resistance was detected. We also excluded SALL1 OE cells in which no Hygromycin-resistance transcripts were detected.

After defining the top highly variable genes (vst method, 5,000, mitochondrial genes and ribosomal genes excluded), data scaling (cell-cycle scores, percentage of mitochondria genes and number of detected genes regressed out) and PCA were performed (30 dimensions). CSS was further applied to integrate data of different samples, followed by PCA on CSS to obtain the PCA-reduced CSS representation, which was then used to generate the UMAP embedding as well as Louvain clustering (resolution = 0.2). Cluster markers were calculated using FindAllMarkers function from Seurat with a Wilcoxon test, thresholding for expression in minimum 25% and logFC > 1.2.

Similarity scores of control and co-cultured MGL to primary microglia were generated by calculating the RSS (simspec package 49), as well as label transfer from primary microglia to MGL were generated as for the co-cultured MGL (see above). For differential gene expression analysis based on the negative binomial distribution between different culture conditions at different time points, pseudobulk samples were generated and analysed using the package DESeq2 58 (v1.38.3) with design = replicate + condition and genes with abs(logFC) > 1.5 and BH-adjusted p-value < 0.01 considered as DE. Pathway enrichment analysis of DEGs between control and co-culture in various cell types was performed with the R package enrichR 85 (v3.4) with default settings and extracting information from the GO Biological Process 2023 database.

### SALL1 OE MGL-organoid co-culture data preprocessing and analysis (droplet-based)

Cell Ranger (10x Genomics, v4.0.0) was used to map the sequencing reads to the human reference (GRCh38-based, 10x Genomics, v3.0.0) and call cells to generate the count matrices. Next, the scRNA-seq data from MGL generated by the two different protocols was merged and normalized using Seurat (v4.0.0)82. Additional quality control was done by filtering out cells with less than 700 genes detected or with mitochondrial transcript percentage higher than 30%.

The scRNA-seq data of all organoid SALL1 OE co-culture samples was merged, followed by normalization, highly variable gene identification (vst method, 5,000, mitochondrial genes and ribosomal genes excluded), data scaling and PCA. CSS was further applied to integrate data of different samples, followed by PCA on CSS to obtain the PCA-reduced CSS representation (with 30 dimensions), which was then used to generate the UMAP embedding as well as Louvain clustering (resolution = 0.2). Cluster markers were calculated using FindAllMarkers function from Seurat with a Wilcoxon test, thresholding for expression in minimum 25% and logFC > 1.2. The resulting clusters were initially annotated as “organoid” or “microglia” and subset into two objects - one microglia and one organoid one, to be analysed separately. The two objects - microglia and organoid - were preprocessed separately (finding top 5,000 variable genes, scaling data, running PCA and integrating with CSS.

For the microglia object and for the organoid object, separately, Louvain clusters with resolution 0.3 or resolution 0.1, respectively, were further annotated based on cluster markers generated with FindAllMarkers function from Seurat with a Wilcoxon test, thresholding for expression in minimum 25% and logFC > 1.2. Similarity scores of control and OE co-cultured MGL to primary microglia were generated by calculating the RSS (simspec package 49).

Differential gene expression between control and co-culture cells was calculated with Wilcoxon Rank Sum test from the presto package 87 (v1.0.0) (p-value < 0.01 and log2 fold change > 0.04) for organoid cells or using the package DESeq2 58 (v1.38.3) and genes with abs(logFC) > 0.18 and BH-adjusted p-value < 0.01 considered as DE for pseudobulk MGL cells. GO of DEGs was performed using the R package enrichR 85 with the database “KEGG 2021 Human” or “GO Biological Process 2025” for organoid or MGL cells, respectively.

Pathway activity inference of MGL cells was performed using the decoupleR package 89 (v2.10.0) with the wrapper to access Pathway RespOnsive GENes for activity inference (PROGENy) model gene weights. Inference of changes in cellular communication analysis of organoid and MGL cells between control and SALL1 OE conditions was performed using the package (single-cell Differential Communication) scDiffCom 72 (v1.1.1). We used the integrated in the package internal collection of ligand-receptor interactions retrieved from seven curated databases.

### SALL1 OE MGL-organoid co-culture data preprocessing and analysis (MARS-seq)

Index-sorted, plate-based singlecell RNA sequencing (scRNA-seq) data were analyzed using the Seurat R package (version 4.3.1). Cells with number of genes < 150 > 4,000, number of transcripts < 11,000, percent.mt > 10%, ribosomal gene content > 6% , and hemoglobin gene > 5% content were excluded. Following these initial filters, additional genes—including those related to mitochondria, ribosomes, hemoglobin, immunoglobulins, and pseudogenes—were removed. Genes with fewer than three total counts across all cells were also excluded. The data were then log-normalized, and highly variable genes were identified using the FindVariableFeatures function. Linear transformation scaling was applied prior to performing principal component analysis (PCA) via the RunPCA function. A nearest-neighbor graph was constructed using Euclidean distances in PCA space (FindNeighbors), and clustering was carried out using the Louvain algorithm implemented in the FindClusters function with default parameters. Single cells were visualized using UMAP or diffusion maps, and differentially expressed markers were identified using FindAllMarkers, applying a log fold change threshold of 0.25 and a minimum expression fraction (min.pct) of 0.1.

## Supplementary tables

**Table S1. Table with abbreviations throughout the text**.

**Table S2. Table with DE genes from DESeq2 analysis of McQuade vs Brownjohn**.

**Table with DE genes between control and 30 day co-cultured MGL**.

**Table S4. Table with genes used from the query dataset for all RSS similarity calculations**.

**Table S5. Table with DE genes between control and 30 day co-cultured forebrain neuron progenitor cells**.

**Table S6. Table with DE genes between control and 30 day co-cultured hindbrain neuron progenitor cells**.

**Table S7. Table with DE genes between control and 30 day co-cultured immature choroid plexus**.

**Table S8. Table with DE genes between GFP control and SALL1 OE MGL co-cultured with organoids for 15 days**.

**Table S9. Overview 10X experiments**.

## Supplementary figures

Starting on page 24.

**Figure S1.**
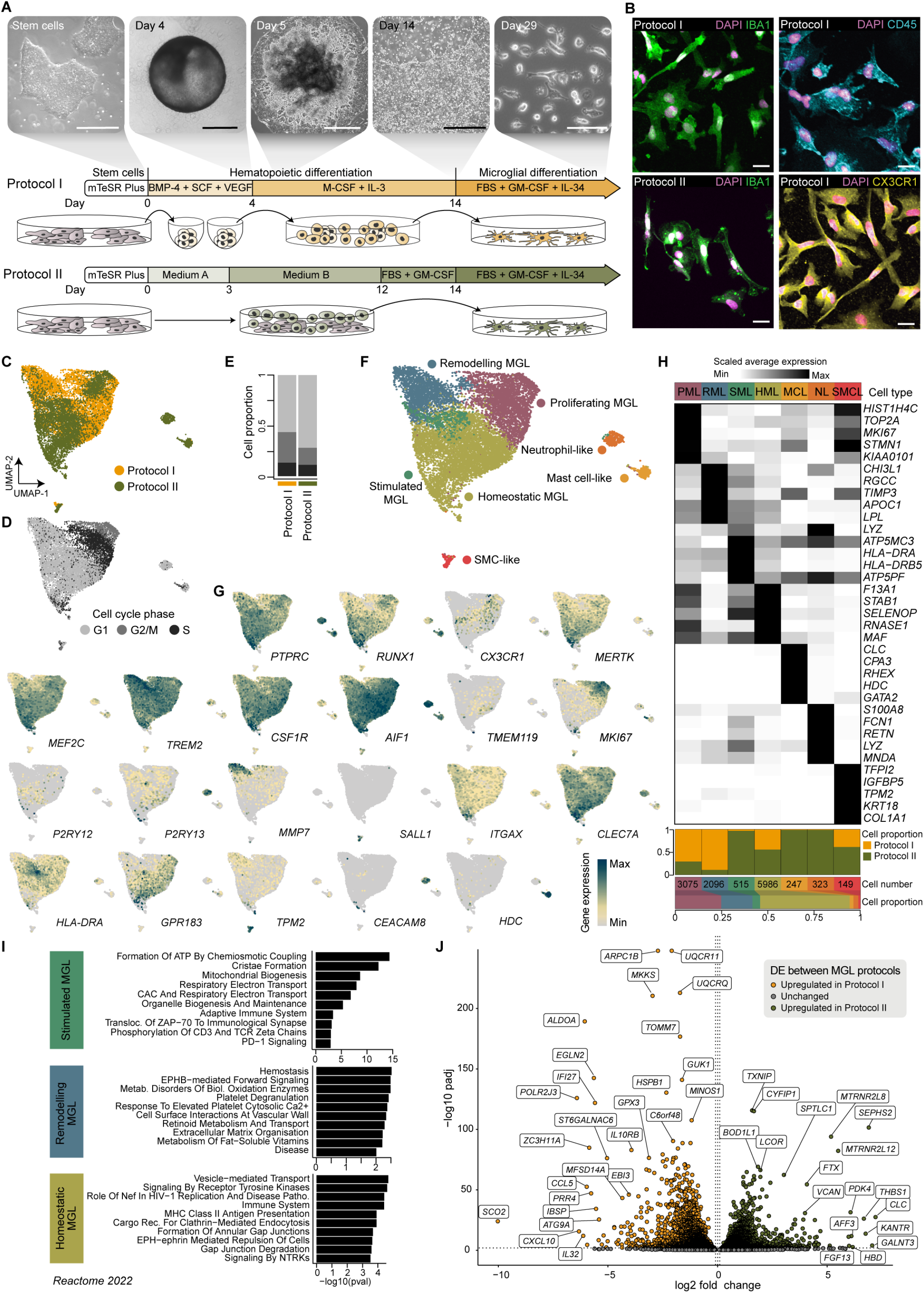
Single-cell transcriptome comparison of Microglia-like cells (MGL) derived from two different protocols. Related to Fig. 1. (A)Schematic representation of the two protocols for generation of stem-cell derived microglia (Protocol I based on ^19^ and ^42^, Protocol II based on ^43^) with brightfield images belonging to key time points in Protocol I. (B) Immunohistochemistry of MGL derived from Protocol I or II marking IBA1 (green) and nuclei (magenta, DAPI) (left) as well as MGL derived from Protocol I marking the tyrosine phosphatase CD45 (cyan, top), the chemokine receptor CX3CR1 (yellow, bottom) and nuclei (DAPI, magenta) (right). MGL from both protocols show comparable morphology and IBA1 expression. (C, D) UMAP representation of MGL colored by Protocol of origin (C) or by cell cycle phase (D). (E) Proportion of cells generated with Protocol I or II in cell cycle phases. (F) UMAP of MGL from Protocol I and II colored by cell type. (G) Feature plots of microglial marker genes and top marker genes of MGL across the two protocols. (H) Heatmap of scaled expression (z-score) of top 5 cell type markers, including proportion of cells from each protocol, number and proportion of cells per cell type. (I) Pathway (Reactome 2022) enrichment analysis of DE genes marking MGL clusters ‘Activated’, ‘Remodeling’ and ‘Homeostatic’. (J) DE genes between MGL derived with Protocol I (yellow) and Protocol II (green). Scale bar: A, left to right - 400 µm, 400 µm, 400 µm, 150 µm, 100 µm; B - 25 µm. BMP-4 - bone morphogenetic protein 4, SCF, VEGF - vascular endothelial growth factor, M-CSF - macrophage colony-stimulating factor, IL-3 - interleukin 3, GM-CSF - granulocyte-macrophage colony-stimulating factor, IL-34 - interleukin 34, FBS - fetal bovine serum, UMAP - Uniform Manifold Approximation and Projection, MGL - induced microglia-like cells, PML - proliferating MGL, RML - remodeling MGL, AML - activated MGL, HML - homeostatic MGL, MCL - mast cell-like, NCL - neutrophil-like, SMCL - smooth muscle cell-like, DE - differentially expressed.

**Figure S2.**
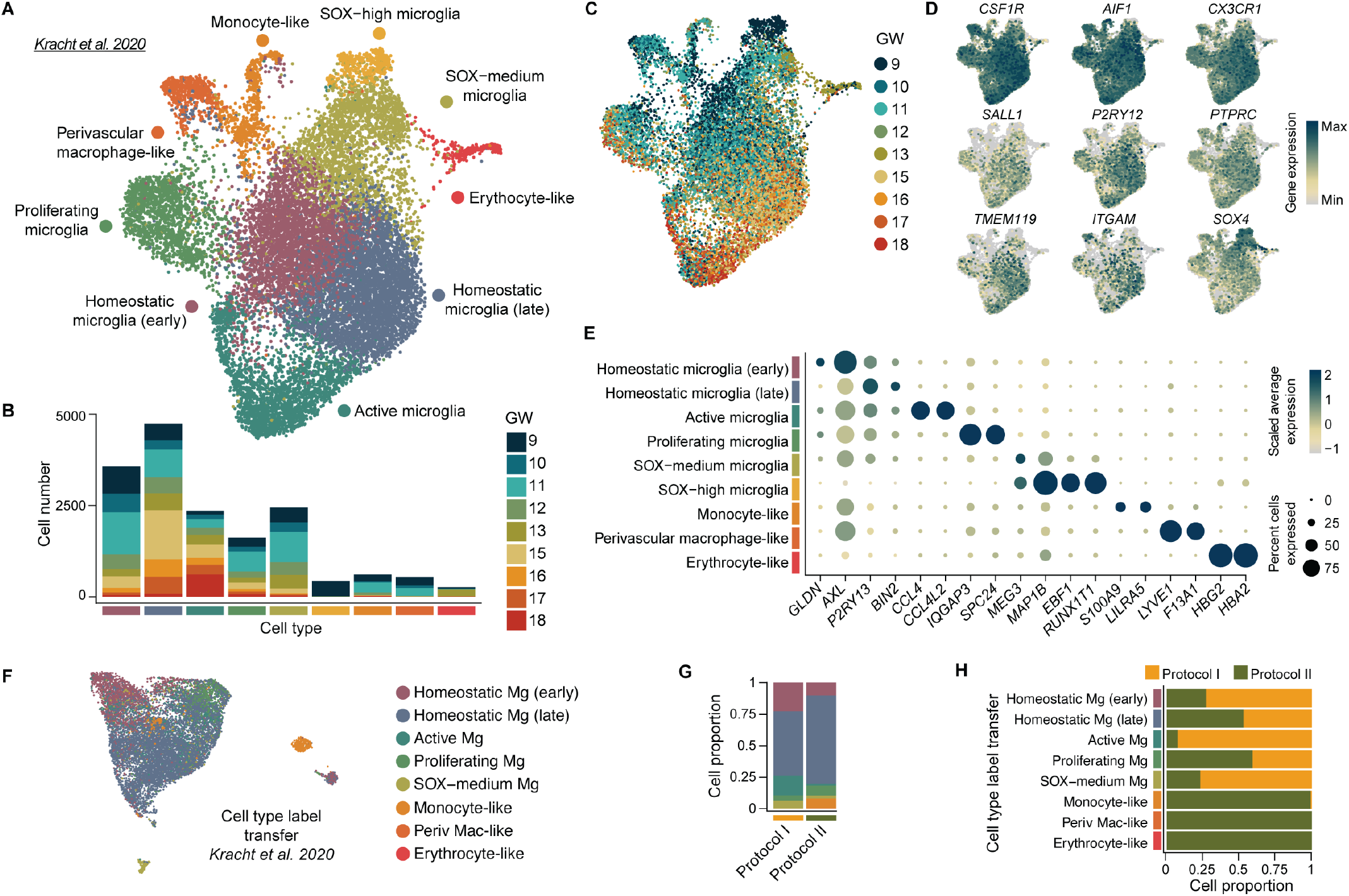
Heterogeneity in human embryonic microglia and reference mapping of Microglia-like cells. Related to Fig. 1, Fig. 3, Fig. 4. (A)UMAP of primary human microglia integrated with Cluster Similarity Spectrum ^49^ and colored by cluster annotated based on analysis performed in this study ^46^. (B) Proportion of cells from each GW contributing to microglia clusters. MGL most closely resemble primary microglia from gestational week 9-15. (C) UMAP of human microglia colored by GW of sample origin. (D) Feature plots of microglial marker genes in primary microglia. (E) Gene expression (color) and proportion of cells per cell cluster (dot size) of marker genes of the primary microglia dataset. (F, G) Label transfer of primary microglia ^46^ onto MGL from in vitro derivation Protocol I and II, presented onto a UMAP (see Figure S1F) (F) and as a stacked barplot indicating proportion of cells per protocol (G). (H) Proportion of cells from each protocol in MGL cell types grouped by label transfer from primary human microglia ^46^. Mg - microglia, GW - gestational week.

**Figure S3.**
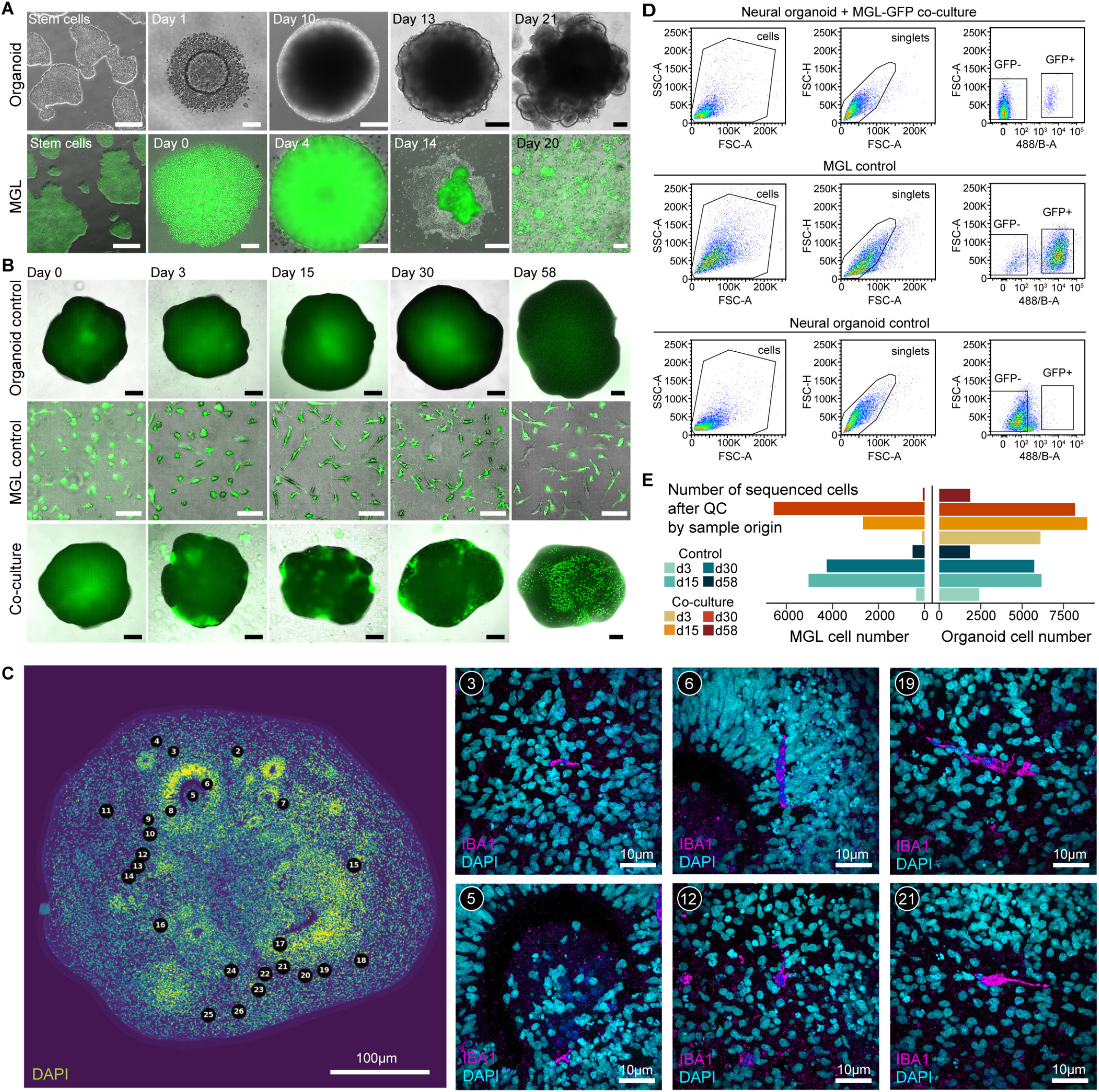
Characterization of organoids and MGL. Related to Fig. 1. (A) Brightfield (organoid) and overlaid brightfield and fluorescence (MGL) images at key steps of differentiation. (B) Overlaid brightfield and fluorescence images at key steps of differentiation of organoid control, MGL control and co-cultured organoids with MGL right before co-culture (day 0) and at day 3, 15, 30 and 58. (C) Most-left panel: Overview of a slice from a fixed 8-week-old neural organoid co-cultured with MGL for 4 weeks with nuclei counterstained with DAPI. Locations of IBA1-positive cells are marked with a location number. Panels on the right: Higher magnification images of selected locations with detected IBA1-positive cells. (D) Exemplary fluorescence activated cell sorting plots of organoids co-cultured with MGL, MGL control and organoid control cells and gating strategy for GFP-positive and -negative cells enrichment. (E) Number of sequenced cells from each sample after QC, used for all further analysis. Scale bar: A, from left to right, both rows - 500 µm, 100 µm, 200 µm, 200 µm, 200 µm; B, organoid control and co-culture images - 500 µm, MGL images - 100 µm. GFP - green fluorescence protein, QC - quality control, d[…] - day […].

**Figure S4.**
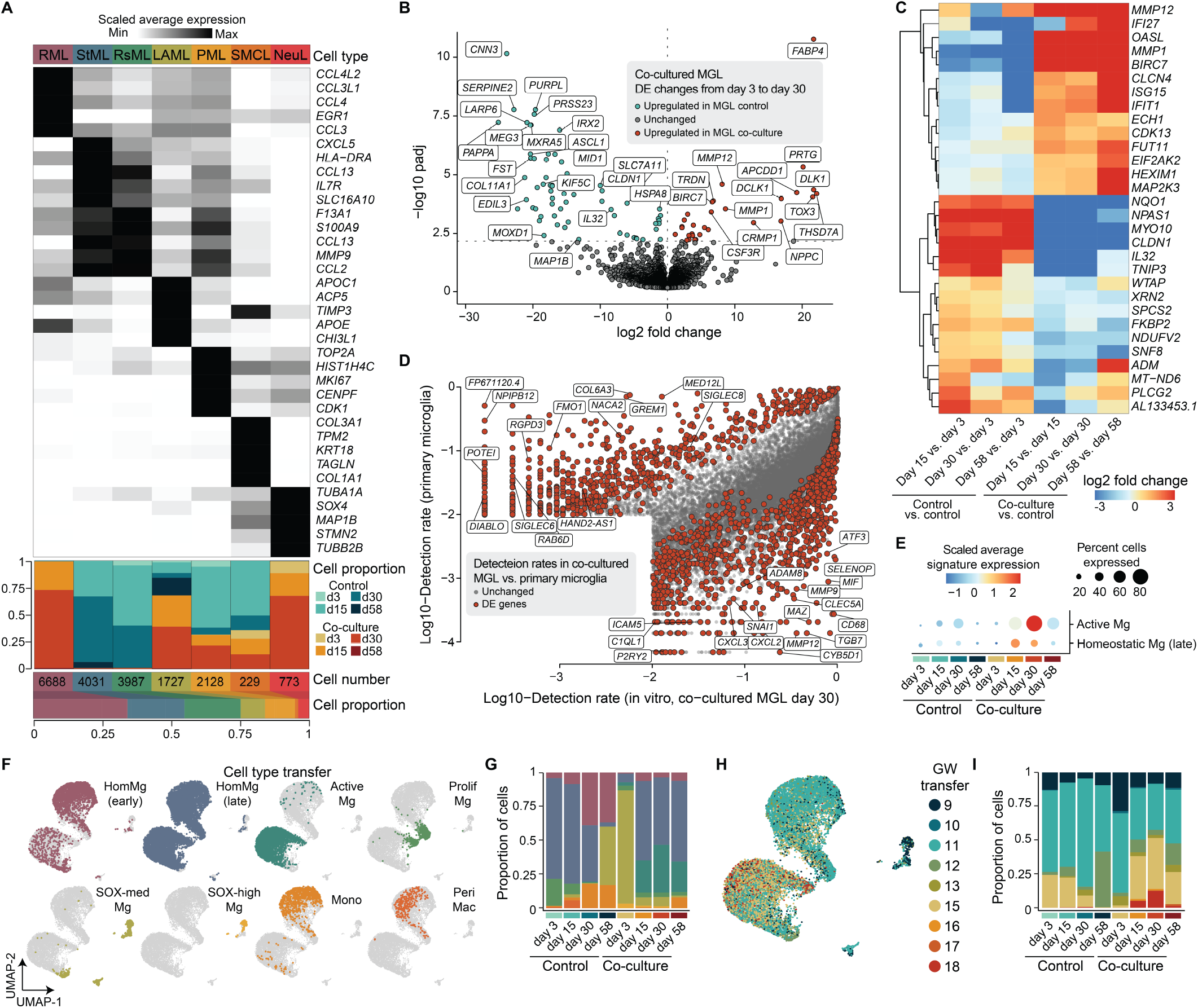
Extended analysis of MGL co-culture within neural organoids. Related to Fig. 1. (A)Heatmap of scaled gene expression (z-score) of top 5 cell type markers with proportion of cells from each sample, number of cells and proportion of cells for each cell type. (B) DE genes from day 3 to day 30 of (co-)culture between control and co-cultured MGL. (C) Heatmap of scaled gene expression changes (z-score) in control MGL and between control and co-cultured MGL as a function of time. (D) Scatter plot of (log10) detection rates of common genes in primary and 30-day co-cultured MGL, with all DE genes colored in red. (E) Scaled average expression (color) of signature genes and percentage of cells expressing those genes (dot size) of ‘Active’ and ‘Homeostatic (late)’ primary microglia across MGL control and co-culture samples. (F, G) UMAP representations (F) and proportion of cells in each sample (G), control and co-cultured MGL, colored by cell type based on label transfer from primary human microglia ^46^. (H, I) UMAP representation (H) and proportion of cells in each sample (I), control and co-cultured MGL, colored by GW based on label transfer from primary human microglia ^46^. RML - remodeling MGL, AML - activated MGL, RsML - resting MGL, LAML - lipid-associated MGL, PML - proliferating MGL , SMCL - smooth muscle cell-like, NeuL - neuron-like, d[…] - day […], GW - gestational week.

**Figure S5.**
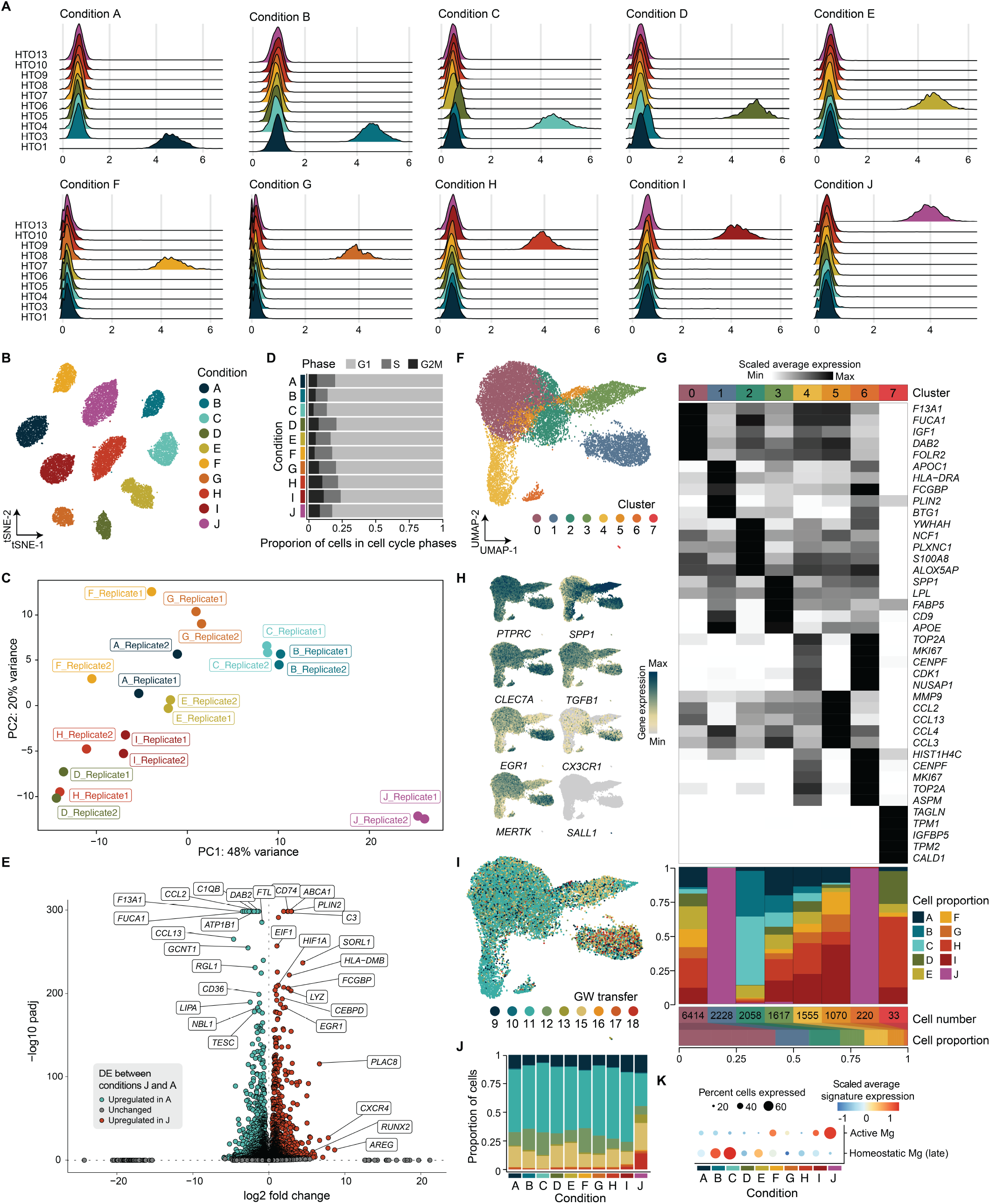
MGL replicate consistency and within treatment heterogeneity. Related to Fig. 3. (A) Ridge plot visualization of HTO enrichment across conditions. (B) t-distributed Stochastic Neighbor Embedding of detected HTOs, excluding doublets and HTO-negative droplets. (C) PC analysis plot of pseudobulk scRNA-seq data from MGL conditions split by biological replicate (‘Replicate1’ and ‘Replicate2’). (D) Proportion of cells in G1, S or G2M cell cycle phases in each MGL culture condition. (E) DE genes between MGL differentiated with media from condition J (cyan) versus those with condition A (red). (F) UMAP of MGL cultured in different media conditions colored based on cluster. (G) Heatmap of scaled gene expression (z-score) of top 5 cluster marker genes across clusters, proportion of cells from each condition belonging to each cluster, number of cells and proportion of cells each cluster contributes to the total cell number. (H) Feature plots of gene expression of microglia markers in MGL treated with cultured media conditions A-J. (I, J) UMAP representation (I) and proportion of cells in each condition (J) colored by GW based on label transfer from primary human microglia ^46^. (K) Scaled average expression (color) of signature genes and percentage of cells expressing those genes (dot size) of ‘Activated’ and ‘Homeostatic (late)’ primary microglia across MGL culture conditions. HTO - hashtag oligo, DE - differentially expressed, PC - principal component, Mg - microglia.

**Figure S6.**
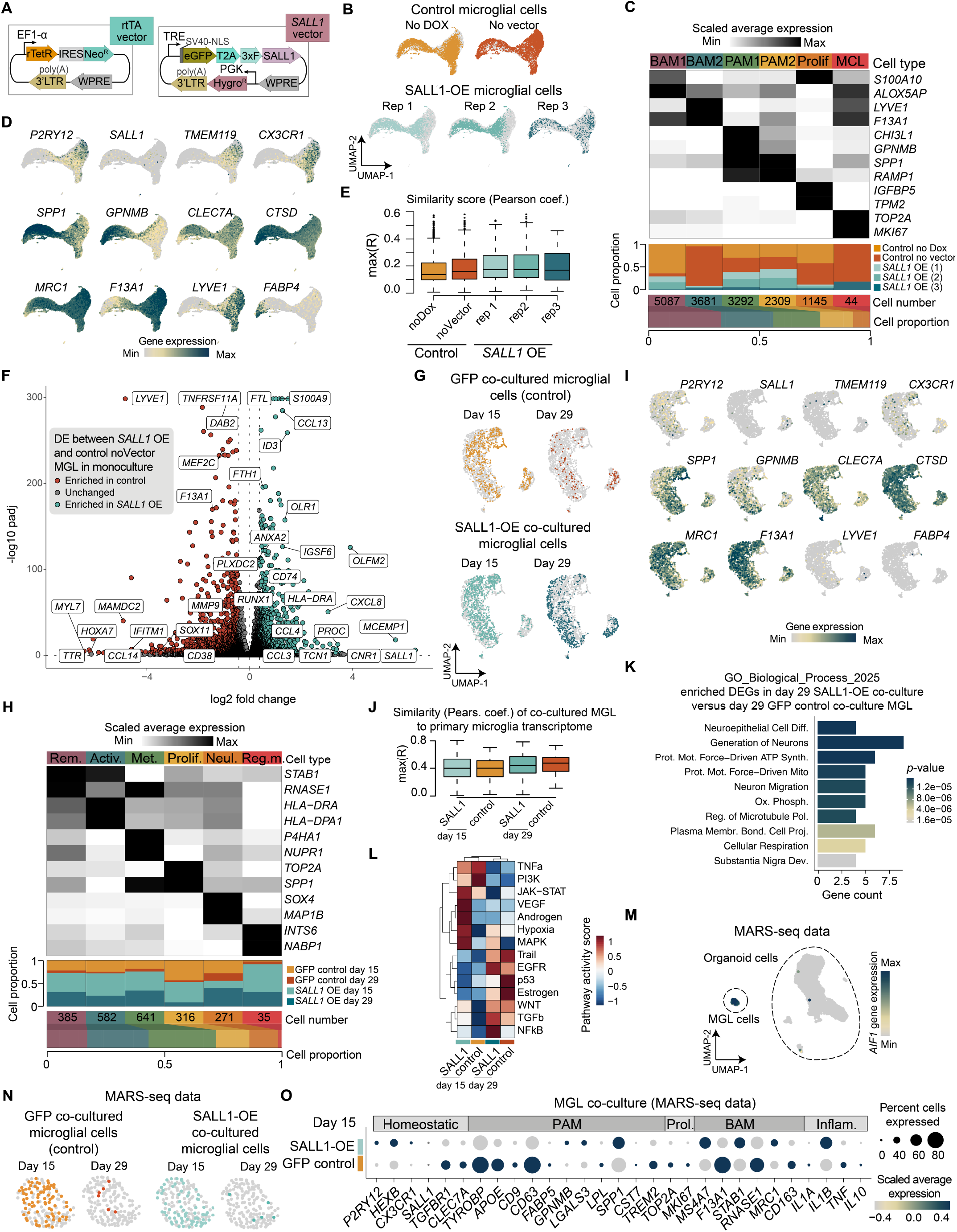
Extended analysis of SALL1 overexpression in MGL co-cultured within neural organoids. Related to Fig. 4. (A)Schematics of the two vectors used for MGL transfection for SALL1 overexpression. The control GFP vector contained only eGFP instead of SALL1 fused to eGFP as shown in the schematic. (B) UMAP of 2D monoculture MGL colored by condition. (C) Heatmap of scaled gene expression (z-score) of top 2 cluster marker genes across cell types, proportion of cells from each condition belonging to each cell type, number of cells and proportion of cells each cell type contributes to the total cell number. (D) Feature plots of homeostatic, proliferative-region-associated microglia and border-associated macrophage marker gene expression in 2D monoculture of MGL. (E) RSS similarity score of 2d monoculture of MGL to meta-cells of a published primary fetal human microglia scRNA-seq dataset ^46^ presented as a boxplot. (F) Volcano plot of differentially expressed genes between SALL1 OE and Control noVector MGL in 2D monoculture. (G) UMAP of MGL co-cultured with neural organoids for 15 or 29 days colored by condition (droplet-based scRNA-seq data). (H) Heatmap of scaled gene expression (z-score) of top 2 cluster marker genes across cell types, proportion of cells from each condition belonging to each cell type, number of cells and proportion of cells each cell type contributes to the total cell number. (I) Feature plots of homeostatic, proliferative-region-associated microglia and border-associated macrophage marker gene expression in GFP control or SALL1 OE MGL co-cultured with neural organoids for 15 or 29 days. (J) RSS similarity score of MGL that has been co-cultured with neural organoids to meta-cells of a published primary fetal human microglia scRNA-seq dataset ^46^ presented as a boxplot. (K) GO analysis of genes upregulated in SALL1 OE MGL in comparison to GFP control MGL in neural organoid co-culture for 29 days. (L) Heatmap presenting pathway activity inference for MGL co-cultured with neural organoids, analysed using PROGENy ^71^. (M) Feature plot of all organoid and MGL cells, sequenced with MARS-seq, showing the expression of AIF1, a macrophage marker. (N) UMAP of MGL co-cultured with neural organoids for 15 or 29 days colored by condition (MARS-seq data). (O) Dotplot showing scaled gene expression (color) and percentage of cells expressing the respective gene (dot size) of homeostatic, PAM, proliferating, BAM or inflammation-related markers for GFP control and SALL1 OE MGL co-cultured with organoids for 15 days (MARS-seq data). BAM - border-associated macrophage, PAM - proliferative-region associated microglia, MCL - muscle-like cell.

**Figure S7.**
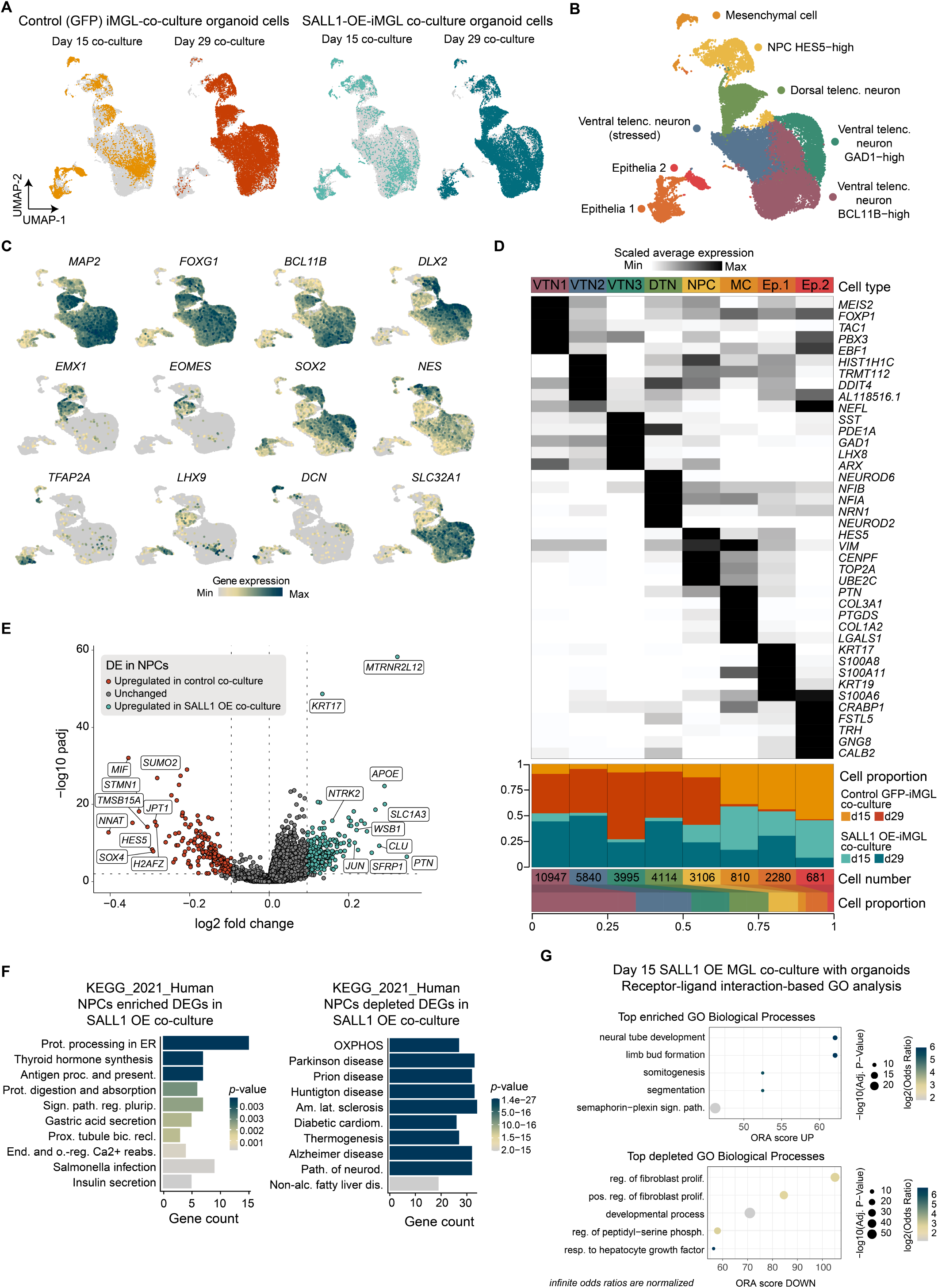
Extended analysis of neural cells in neural organoids containing SALL1-overexpressed MGL. Related to Fig. 4. (A, B) UMAP of scRNA-seq data of neural organoid cells from co-cultures with GFP control or SALL1 OE MGL, colored by condition (A) or by cell type (B). (C) Feature plots of marker genes in organoids. (D) Heatmap of scaled gene expression (z-score) of top 5 cluster marker genes across cell types, proportion of cells from each condition belonging to each cell type, number of cells and proportion of cells each cell type contributes to the total cell number. (E) Volcano plot of differentially expressed genes between NPCs from organoids co-cultured with SALL1 OE and GFP control MGL. (F) KEGG pathway enrichment analysis of processes mapping to genes enriched (left) or depleted (right) in the NPCs of organoids co-cultured with SALL1 OE MGL in comparison to co-cultures with control MGL. (G) GO for biological processes enriched (top) or depleted (bottom) in co-cultures of organoids with SALL1 OE MGL in comparison to co-cultures with control MGL, based on receptor-ligand interaction analysis using scDiffCom ^72^. VTN1 - ventral telencephalic neuron BCL11B-high, VTN2 - ventral telencephalic neuron (stressed), VTN3 - ventral telencephalic neuron GAD1-high, DTN - dorsal telencephalic neuron, NPC - neural progenitor cell, MC -mesenchymal cell, Ep. - epithelial cells, DE - differential expression, KEGG - Kyoto Encyclopedia of Genes and Genomes, GO - gene ontology, ORA - over-representation analysis.

